# Minnesota peat viromes reveal terrestrial and aquatic niche partitioning for local and global viral populations

**DOI:** 10.1101/2020.12.15.422944

**Authors:** Anneliek M. ter Horst, Christian Santos-Medellín, Jackson W. Sorensen, Laura A. Zinke, Rachel M. Wilson, Eric R. Johnston, Gareth G. Trubl, Jennifer Pett-Ridge, Steven J. Blazewicz, Paul J. Hanson, Jeffrey P. Chanton, Christopher W. Schadt, Joel E. Kostka, Joanne B. Emerson

## Abstract

**Background:** Peatlands are expected to experience sustained yet fluctuating higher temperatures due to climate change, leading to increased microbial activity and greenhouse gas emissions. Despite mounting evidence for viral contributions to these processes in peatlands underlain with permafrost, little is known about viruses in other peatlands. More generally, soil viral biogeography and its potential drivers are poorly understood at both local and global scales. Here, 87 metagenomes and five viral size-fraction metagenomes (viromes) from a boreal peatland in northern Minnesota (the SPRUCE whole-ecosystem warming experiment and surrounding bog) were analyzed for dsDNA viral community ecological patterns, and the recovered viral populations (vOTUs) were compared to our curated PIGEON database of 266,805 vOTUs from diverse ecosystems.

**Results:** Within the SPRUCE experiment, viral community composition was significantly correlated with peat depth, water content, and carbon chemistry, including CH_4_ and CO_2_ concentrations, but not with temperature during the first two years of warming treatments. Peat vOTUs with aquatic-like signatures (shared predicted protein content with marine and/or freshwater vOTUs) were significantly enriched in more waterlogged surface peat depths. Predicted host ranges for SPRUCE vOTUs were relatively narrow, generally within a single bacterial genus. Of the 4,326 SPRUCE vOTUs, 164 were previously detected in other soils, mostly peatlands. None of the previously identified 202,372 marine and freshwater vOTUs in our PIGEON database were detected in SPRUCE peat, but 1.9% of 78,203 genus-level viral clusters (VCs) were shared between soil and aquatic environments. On a per-sample basis, vOTU recovery was 32 times higher from viromes compared to total metagenomes.

**Conclusions:** Results suggest strong viral “species” boundaries between terrestrial and aquatic ecosystems and to some extent between peat and other soils, with differences less pronounced at the “genus” level. The significant enrichment of aquatic-like vOTUs in more waterlogged peat suggests that viruses may also exhibit niche partitioning on more local scales. These patterns are presumably driven in part by host ecology, consistent with the predicted narrow host ranges. Although more samples and increased sequencing depth improved vOTU recovery from total metagenomes, the substantially higher per-sample vOTU recovery after viral particle enrichment highlights the utility of soil viromics.

## Background

Peatlands store approximately one-third of the world’s soil carbon (C) and have a significant role in the global C cycle [1]. Microbial activity in peatlands plays a key role in soil C and nutrient cycling, including soil organic C mineralization to the greenhouse gases, methane (CH_4_) and carbon dioxide (CO_2_) [2–5]. Given the abundance of viruses in soil (10^7^ to 10^10^ per gram of soil [6–9]) and evidence for viral impacts on microbial ecology and biogeochemistry in other ecosystems [10–12], it is likely that viral infection of soil microorganisms influences the biogeochemical and C cycling processes of their hosts [13–15]. In marine ecosystems, viruses are estimated to lyse 20-40% of ocean microbial cells daily, impacting global ocean food webs and the marine C cycle [16–18], and viral contributions to terrestrial ecosystems are presumed to be similarly important but are less well understood [6,13,14,19–21].

Our current understanding of soil viral ecology stems from pioneering studies on viral abundance, morphology, amplicon sequencing, and lysogeny of bacteria [22–27], along with early viral size-fraction metagenomic (viromic) investigations [28–30]. More recently, total soil and wetland metagenomic datasets have been mined for viral sequences in a subarctic peatland spanning a natural permafrost thaw gradient [15], a freshwater marsh [10], and through a global meta-analysis [31], revealing thousands of previously unknown viral populations (vOTUs) and suggesting habitat specificity for some of these viruses. An effort to mine metatranscriptomic data for RNA viruses in Mediterranean grasslands revealed differences in RNA viral communities in bulk, rhizosphere, and detritusphere (plant litter-influenced) soil compartments [32]. Similar mining of metatranscriptomic data from peat bog *Sphagnum* mosses revealed that viruses may play an important role in the ecology of the *Sphagnum* microbiome [33]. In addition to mining omic data for viral signatures, laboratory enrichment of viral particles prior to sequencing can allow for the generation and analysis of viral size-fraction metagenomes (viromes) from soil. This approach has recently been paired with high-throughput sequencing, revealing more comprehensive insights into soil viral ecology [13,15,34,35], including in thawing permafrost peatlands.

Thawing permafrost peatlands have been the focus of several recent viral and other microbial diversity studies that seek to better understand ecological patterns underlying C emissions from these climate-vulnerable ecosystems [13,15,36–38]. Microbial (bacterial and archaeal) diversity tends to be highest in surface peat and decreases with depth [1,38–42], and similarly, viral community composition has been shown to vary by depth in the seasonally thawed active layer of permafrost [15]. These peat soils were characterized by relatively high viral diversity (thousands of vOTUs), including viruses predicted to infect methanogens and methanotrophs that are responsible for CH_4_ cycling [15]. Viruses and other microbes have been shown to be active in the active layer of permafrost through metatranscriptomics, and bacterial and/or archaeal activity has also been shown through stable isotope probing and metaproteomics [15,38,43,44]. Furthermore, both microbial and viral community composition have been shown to differ according to permafrost thaw stage, suggesting that these microbiota and their coupled dynamics could change with changing climate [13,15,36,38]. Evidence for more direct viral impacts on ecosystem C cycling has been revealed by the recovery of putative viral auxiliary metabolic genes (AMGs) [13,15], specifically, virus-encoded glycosyl hydrolases capable of degrading complex C into simple sugars [15].

Although we are gaining insights into soil viral ecology within specific ecosystems, global soil viral biogeographical patterns and their underlying drivers are largely unknown. Cultivation- and genomics-based studies of mycobacteriophages have revealed that some closely related phage isolates and genome clusters are widespread across the globe, while others seem to be more geographically restricted, often contained in a single region of the USA, where the majority of the samples were collected [38,39]. A study of T4-like g23 major capsid gene amplicons in rice paddy floodwater revealed that T4-like phage communities changed with sampling time and location and that these communities were mostly structured by geographical separation, but also by ecological environment (*e.g*., freshwater, soil, marine, or wetland environments), such that the phage communities were more similar in the same ecological environments [27]. In better studied marine ecosystems, virions are thought to be transported along oceanic currents and by sinking particles, and viral communities tend to be structured locally by environmental factors that affect host microbial communities [45]. In general, marine viruses occupying similar habitats tend to be closely related, in terms of shared sequence homology and community composition, even across large geographic distances [31]. However, given the substantial physicochemical differences between relatively well-mixed marine and highly heterogeneous, structured terrestrial ecosystems [14], together with the relative dearth of information on soil viruses, it is difficult to predict the extent to which previously identified biogeographical patterns in marine viral communities might also apply to soil.

Although we now have an array of laboratory and bioinformatics methods for soil viral ecology [7,15,23,31,34,46–51], we lack a thorough comparative understanding of these approaches and best practices. As one specific example, viral size-fraction metagenomes (viromes) from grassland and agricultural soils have been shown to be substantially enriched in viral and ultrasmall cellular organismal DNA, compared to total metagenomes that tend to be more enriched in DNA from cellular organisms too large to easily pass through the 0.2 μm filters used for viral enrichment [35,52]. Although these results would suggest that viromes may be more appropriate than total metagenomes for studying viral communities, the generalizability of this trend across soils and in other ecosystems is unknown. In fact, a recent meta-analysis of human gut sequencing data reported that total metagenomes may recover more viral sequences than viromes [53], though the available datasets for that analysis generally precluded robust, direct comparisons of both approaches applied to the same samples.

In this study, we examined viral communities in boreal peatlands in Minnesota, USA. Cold, acidic, and waterlogged conditions in these peatlands slow decomposition, resulting in C accumulation over centuries [54]. Rising temperatures, changing hydrology, and oxygenation of surface peats are predicted to accelerate decomposition of the accumulated C, increasing ecosystem respiration and enhancing greenhouse gas emissions as a positive feedback to climate change [54,55]. The Marcell Experimental Forest (MEF) in Minnesota, USA is at the southern edge of the boreal zone and is expected to be particularly vulnerable to climate change [54]. MEF has been the site of numerous studies on greenhouse gas emissions, C sequestration, hydrology, biogeochemistry, and vegetation [56–61]. To investigate the response of peatlands to increasing temperature and atmospheric CO_2_ concentrations, the US Department of Energy (DOE) established the Spruce and Peatland Responses Under Changing Environments (SPRUCE) experiment in MEF. This experiment is within an intact peat bog ecosystem, consisting of *Picea mariana* (black spruce) and *Larix laricina* (larch) trees, an ericaceous shrub layer, and a predominant cover of *Sphagnum* with minor contributions of other mosses [54,55,62]. SPRUCE researchers are studying whole-ecosystem responses to temperature and elevated CO_2_ (eCO_2_), including the responses of plants, above- and belowground microbial communities, and whole-ecosystem processes, such as greenhouse gas emissions [1,54,55,63–67].

Here, we used a combination of total soil metagenomics and viromics to: 1) investigate peat viral community composition and its potential drivers in the SPRUCE experiment, 2) place the recovered vOTUs in global biogeographical and ecosystem context, and 3) compare the two approaches (total metagenomics and viromics) for recovering soil viral population sequences. We are also contributing a new database for reference-based viral genome recovery: the <u>P</u>hages and <u>I</u>ntegrated <u>G</u>enomes <u>E</u>ncapsidated <u>O</u>r <u>N</u>ot (PIGEON) database of 266,805 vOTU sequences from diverse ecosystems.

## Results and Discussion

### Dataset overview and peat viral population (vOTU) recovery

To improve our understanding of peat viral diversity, we leveraged 82 peat metagenomes from cores collected from the SPRUCE experiment in northern Minnesota, USA in 2015 and 2016, along with five paired viromes and metagenomes that we collected along a transect outside the experimental plots from the same bog in 2018 at near-surface (top 10 cm) depths. In the field experiment, deep peat heating (DPH) and whole ecosystem warming (WEW) treatments heated the peat (to a depth of 2 m) and air inside chambered enclosures to target temperatures of +2.25, +4.5, +6.75 and +9 °C above ambient temperature [1,55,62,68] inside 8 experimental chambers. There were also two ambient experimental chambers and two unchambered ambient plots (Table S1). Peat samples for metagenomics were collected from four depths (10-20 cm, 40-50 cm, 100-125 cm and 150-175 cm) per year in each chamber and unchambered ambient plot (38 and 44 total soil metagenomes were successfully sequenced in 2015 and 2016, respectively), with approximate sequencing depths of 6 Gbp per metagenome in 2015 and 15 Gbp in 2016. From each of the five transect peat samples (Supplementary Figure 1), a viral size-fraction metagenome (virome) and total soil metagenome were sequenced, each to a depth of approximately 14 Gbp.

Reads from the SPRUCE experiment metagenomes (82), transect viromes (5), and transect total soil metagenomes (5) were assembled into contigs ≥ 10 kbp in length, from which viral contigs were identified [48,49] and clustered into 5,006 approximately species-level viral populations (viral operational taxonomic units, vOTUs [69]). These vOTUs were then clustered with 261,799 vOTUs from diverse habitats in our PIGEON database (see methods, Table S2, available on Dryad (https://datadryad.org/, by DOI of this paper) [10,13,15,31,33,54–58]. The resulting clustered database of 266,805 “species-level” vOTUs from SPRUCE and other ecosystems was then used as a reference for read mapping from each of our metagenomes to identify vOTUs recovered from these peatlands. In total, we recovered 4,326 vOTUs (detected through read mapping) from the SPRUCE experiment and adjacent peatlands. Henceforth, “SPRUCE” refers to our data from the SPRUCE experiment and/or transect, unless otherwise specified.

### Investigating patterns and potential drivers of peat viral community composition in the SPRUCE experimental plots

To characterize peat viral community compositional patterns and their potential drivers, vOTU abundances from the 82 SPRUCE experiment metagenomes were compared to environmental measurements. Using the 4,326 SPRUCE vOTUs as references, we recovered 2,699 vOTUs from the SPRUCE experimental plots through read recruitment and tracked their abundances (average per bp coverage depth) across the experimental plot metagenomes. No significant differences in viral community composition were detected according to temperature treatment (Mantel ρ = 0.0057, p = 0.56), as discussed in more detail below. Viral community composition was significantly correlated with depth (Fig. 1A), even across different temperature treatments and years (Mantel ρ = 0.57, p=0.00001), consistent with previous evidence that viral community composition varies with depth in Swedish peatlands [15] and other soils [70]. These results are also consistent with observations of microbial communities in SPRUCE peat, where depth was shown to explain the largest amount of variation in peat microbial community composition, and temperature effects have thus far (from 2015-2018) been shown not to be significant [1,65]. We also measured a significant difference in viral community composition between the two sampling years (June 2015 and June 2016, PERMANOVA p=0.009), indicating temporal dynamics on time scales shorter than one year. Other factors that significantly (p < 0.05) correlated with viral community composition included microbial community composition, porewater CO_2_ and CH_4_ concentrations, and the calculated fractionation factor for carbon in porewater δ^13^CH_4_ relative to δ^13^CO_2_ (αC) [71] (Table S3), which can be used to infer CH_4_ production and consumption pathways, including whether acetoclastic or hydrogenotrophic methanogenesis is the more dominant pathway [3,15,71,72]. Although all of these factors also co-varied with depth, interestingly, viral community composition was more significantly correlated with αC and porewater CH_4_ concentrations than with depth. Together, these results prompted further exploration of potential explanations for these compositional patterns with depth, including links between SPRUCE vOTUs and water content, peat C cycling, and microbial hosts.

**Figure 1:**
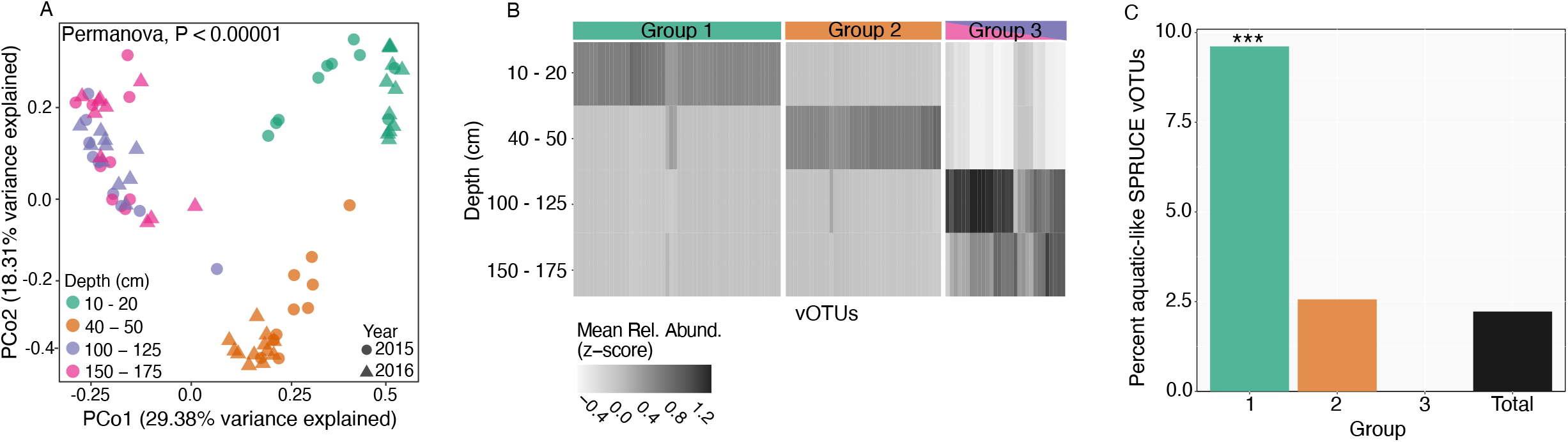
Peat viral community and population (vOTU) abundance patterns with depth in the SPRUCE experimental plots. **A**: Principal coordinates analysis (PCoA) of viral community composition in 82 samples (total soil metagenomes) from peat bog soil from the Marcell Experimental Forest in northern Minnesota (USA) collected from the SPRUCE experimental plots and chambers (temperature treatments ranging from ambient to +9 °C above ambient), based on Bray-Curtis dissimilarities derived from the table of vOTU abundances (read mapping to vOTUs, n=2,699). Each point is one sample (n=82)**. B**: Mean relative abundances (Z-transformed) of vOTUs significantly differentially abundant by depth (adjusted-p<0.05, Likelihood Ratio Test). Groups were identified through hierarchical clustering and are colored according to the depths in panel A. **C**: Percentage of vOTUs classified as “aquatic-like” in each of the groups identified in panel B (Groups 1-3) and in the whole dataset of 2,699 vOTUs (Total). SPRUCE vOTUs were considered “aquatic-like” if they shared a genus-level viral cluster (VC) with at least one vOTU from a marine or freshwater habitat in the PIGEON database. Note that the y-axis maximum is 10%. *** denotes a significantly larger proportion of aquatic-like vOTUs in that group, relative to the proportion of aquatic-like vOTUs in the full SPRUCE dataset (Total) (P < 0.05, Hypergeometric test)

To investigate potential drivers of viral community compositional patterns with depth, we identified 121 vOTUs that exhibited significant differential abundance patterns across peat depth levels (adjusted-p < 0.05, Likelihood Ratio Test). We assigned these vOTUs to one of three groups via hierarchical clustering (Fig. 1B): vOTUs abundant in the near-surface (10-20 cm) but depleted at all other depths, vOTUs abundant in the 40-50 cm depth range but depleted at other depths, and vOTUs abundant in only the two deepest depth ranges (100-125 and 150-175 cm). Given that near-surface peat had significantly higher gravimetric soil moisture measurements than deeper peat (p=0.002, Student’s T-test), and because peat viral community composition was significantly correlated with both depth and measured soil moisture content (Table S3), we investigated the depth-resolved abundance patterns of “aquatic-like” SPRUCE vOTUs. We defined aquatic-like vOTUs as those found in the same “genus-level” viral clusters (VCs) as vOTUs from freshwater and/or marine environments, based on clustering the predicted protein contents of SPRUCE vOTUs with those of aquatic vOTUs in our PIGEON database. Next, we compared the proportion of aquatic-like vOTUs within each of the three depth-range groups and found that the near-surface peat group displayed the highest proportion of aquatic-like vOTUs, followed by the mid-depth group, while the deepest peat group had zero recognizable aquatic-like vOTUs (Fig. 1C). The proportion of aquatic-like vOTUs in the near-surface group deviated significantly from the aquatic-like proportion of the total set of 2,699 vOTUs (p < 0.05, Hypergeometric Test), indicating a significant enrichment of aquatic-like vOTUs in the near surface. Overall, these results suggest that the aquatic-like SPRUCE vOTUs found in the surface horizons and/or their hosts were better adapted to near-surface depths, perhaps due to better adaptation to water-rich environments. Consistent with this interpretation and as is typical for peat sampling, we did not exclude porewater from our samples [3,7,15,37], so it is likely that some of the vOTUs were derived from the porewater directly. Also, although the gravimetric soil moisture content measurements may not accurately reflect peat saturation with depth (water table depth measurements indicated that the entire sampled peat column was saturated for each of the samples), qualitatively, there was substantially more volumetric water content (waterlogging) in the near-surface depths compared to the deeper, more compacted peat. Still, the underlying explanation for the observed enrichment of aquatic-like vOTUs in the near surface could be due to a variety of ecological similarities between near-surface peatlands and aqueous systems beyond simply water content (*e.g*., redox chemistry, substrates, and dissolved oxygen content [36,73]) and warrants further exploration in the future.

Under the assumption that patterns in viral community composition were at least partially indirect, resulting from interactions with hosts, we attempted to bioinformatically link SPRUCE vOTUs to microbial host populations [15]. All 4,326 vOTUs and a total of 486 metagenome-assembled genomes (MAGs), 443 from the SPRUCE experiment metagenomes (Table S4) and 43 from the transect (>60% complete, <10% contaminated, Table S5), were considered in this analysis. A total of 2,870 CRISPR arrays were recovered from the metagenomes via Crass [74], and 29 CRISPR-derived virus-host linkages were made between 23 vOTUs and 21 host MAGs (Fig. 2, Table S6). All 21 of the MAGs were bacterial and could be taxonomically classified to at least the family level, and for each of the six vOTUs linked to more than one host, the predicted hosts were all in the same family. Where genus-level host classification was possible, all vOTUs were predicted to infect the same host genus. However, two vOTUs that were linked to multiple host MAGs had at least one predicted host that could not be classified to the genus level. These results are generally consistent with the expected narrow host range for most viruses, but the data do not exclude the possibility that some of the vOTUs could infect different genera. Of the seven hosts that were predicted to be infected by more than one virus, only one, Acidobacteria bacterium UBA7540 Bin 12, was predicted to be infected by two viruses from the same genus-level viral cluster (VC), meaning that most vOTUs predicted to infect the same host came from different viral genera.

**Figure 2:**
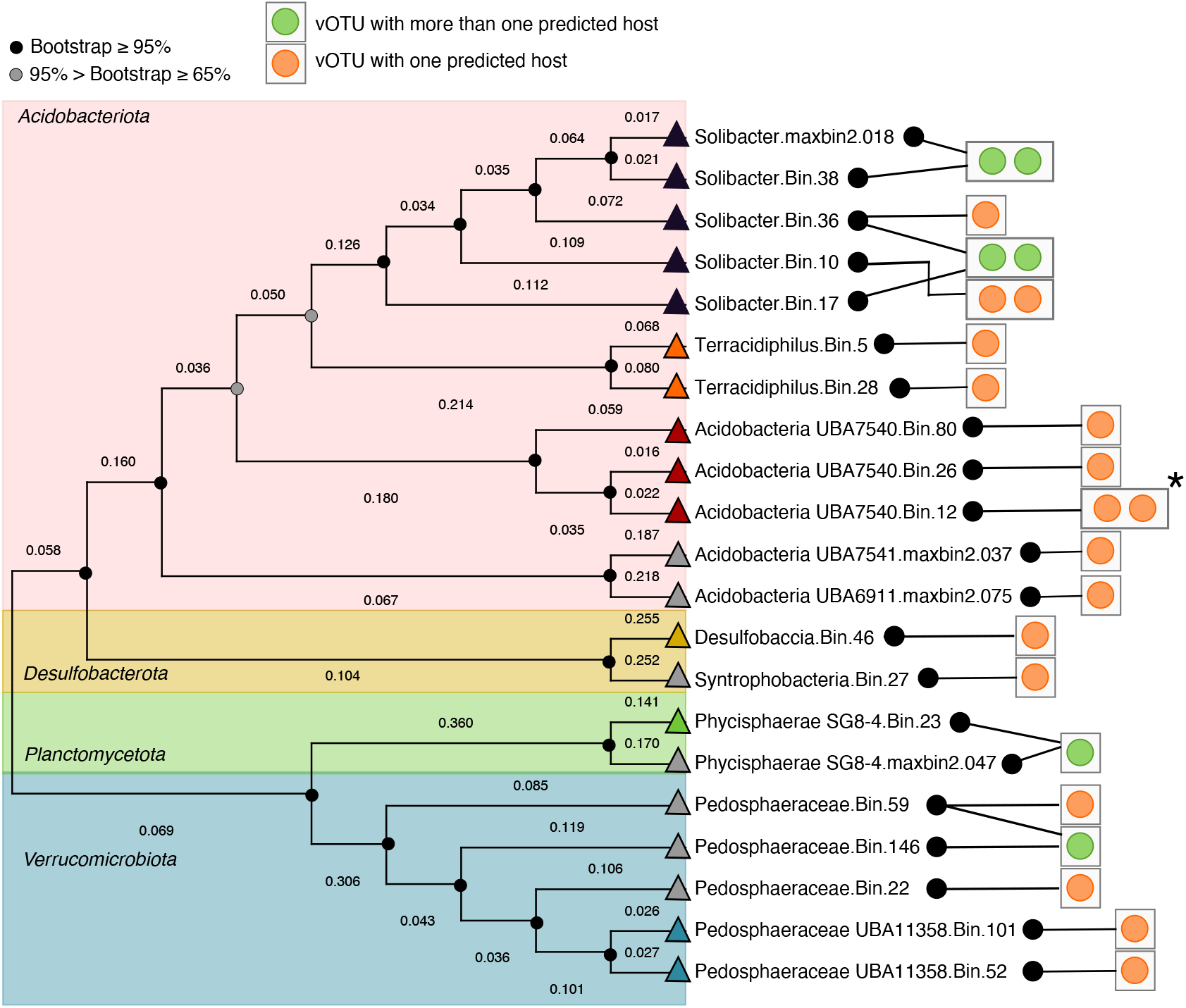
SPRUCE virus-host linkages according to host phylogeny. Unrooted phylogenetic tree (concatenated predicted protein alignment of 43 marker genes defined by CheckM [109]) of microbial host metagenome-assembled genomes (MAGs) with at least one vOTU (green and orange circles) linked via CRISPR sequence homology. Branch lengths represent the expected number of substitutions per site. Lines between black circles and squares with orange or green circles link vOTUs to predicted host MAGs. Colored triangles indicate the MAG genus (the same color is the same genus, except for grey triangles, for which the corresponding MAG could only be classified to the family level). Asterisk indicates vOTUs in the same genus-level viral cluster (VC); remaining vOTUs were all in distinct VCs. Bootstrap support values are shown as circles on nodes, black circles indicate support >= 95%, grey indicates support between 65 and 95%.

To investigate potential connections between virus-host dynamics and environmental conditions, along with viral community links to carbon chemistry, we attempted to assess virus-host abundance ratios and their patterns across samples, and we explored the auxiliary metabolic gene (AMG) content of the vOTUs. Only 10 virus-host pairs (10 vOTUs linked to 9 MAGs) were identified for which both the vOTU and the MAG were detected together in at least one sample, so, unsurprisingly for the small dataset size, significant patterns in virus-host abundance were not found according to any of the parameters considered, including depth, year, αC, CH_4_ and CO_2_ concentrations, and moisture content. To further investigate the significant correlation between αC and viral community composition, we also looked for vOTU linkages to methanogen or methanotroph MAGs, this time based on MAG genomic content as opposed to the above analyses according to MAG taxonomy. HMM searches for McrA (a methanogenesis biomarker) [75,76], sMMO, pMMO, and pXMO (methanotrophy biomarkers) [3] predicted proteins were performed on the 443 SPRUCE experiment MAGs. Nine MAGs were found to contain McrA-encoding genes, and evidence for methanotrophy was found in 22 MAGs, but none of these MAGs had a CRISPR linkage to a vOTU. Thus, we infer either that αC co-varies with an unmeasured variable that better explains viral community composition and/or that important virus-host linkages associated with CH_4_ cycling were not identified through these approaches. Finally, consistent with potential viral roles in the soil C cycle, we identified 287 putative AMGs encoded by viral genomes and predicted to be involved in 18 C-cycling processes, based on VIBRANT output [50] (Supplementary discussion table S7, S8, S9). These results are consistent with previously identified glycosyl hydrolase genes encoded in peat viral genomes [13,15], along with other putative C-cycling AMGs from soil [77,78] (see Supplementary Discussion).

As indicated above, no significant influence of temperature on viral community composition was detected over the first two years of experimental warming. Consistent with these findings, no differences in microbial community composition were found according to temperature treatments in these samples over the first five years of whole ecosystem warming, although warming exponentially increased CH_4_ emissions and enhanced CH_4_ production rates throughout the entire soil profile [65]. These results are also consistent with prior studies that have shown that soil microbial community responses to similar temperature increases can take multiple years to manifest [79–81]. For example, significant differences in soil microbial community composition were found in Harvard Forest after 20 years of soil warming at 5 °C above ambient temperatures [79], after seven years in Austrian forest soils warmed 4 °C above ambient temperatures [80], and after five years of warming the soil only 1.5 °C above ambient temperatures in a *Castanopsis hystrix* plantation (planted forest) [81]. Warming has been shown to substantially alter the community composition, diversity, and N2 fixation activity of peat moss microbiomes [66], and in microcosms of surface peat collected from the SPRUCE site, microbial diversity was negatively correlated with temperature, suggesting that prolonged exposure of the peatland ecosystem to elevated temperatures will lead to a loss in microbial diversity [82]. In the SPRUCE experiment, the fractional cover of *Sphagnum* mosses (*S. magellanicum* and *S. angustifolium/fallax*) decreased with increasing temperature, and the fraction of ground area with no live *Sphagnum* increased with increasing temperature [54]. Plant phenology (the timing of different traits throughout the growing season) also changed for some native plant species [62]. Though no significant temperature response has been observed in the *in situ* belowground peat viral and microbial communities after two to five years of warming, context from these other studies suggests that differences in viral and microbial community composition may follow after a longer period of warming. In addition, evidence for an increased CO_2_ pulse in response to elevated atmospheric CO_2_ concentrations (a manipulation that commenced at SPRUCE after the samples considered here were collected) in combination with warming [65] suggests that changes in belowground communities may also be more readily observed after warming in combination with elevated atmospheric CO_2_ concentrations.

### Placing SPRUCE peat viral “species” in global context

Of the 4,326 vOTUs from SPRUCE, 4,162 were assembled from SPRUCE-associated metagenomes (including the viromes), and 164 were recovered through read mapping to our PIGEON database of vOTUs from diverse ecosystems (Fig. 3A). The previously recovered vOTUs were first reported from other globally distributed sites, mainly peatlands (160 of 164), including peat vOTUs from Sweden (147), Germany (5), Alaska, USA (4), Wisconsin, USA (2), and Canada (2) (Fig. 3B). The recovery of hundreds of viral species (4% of the dataset) in geographically distant peatlands suggests that there may be a peat-specific niche for these viruses. In addition, four vOTUs recovered from SPRUCE peat were first identified in a wet tropical soil in Puerto Rico, suggesting some global species-level sequence conservation across soil habitats (Table S10).

**Figure 3:**
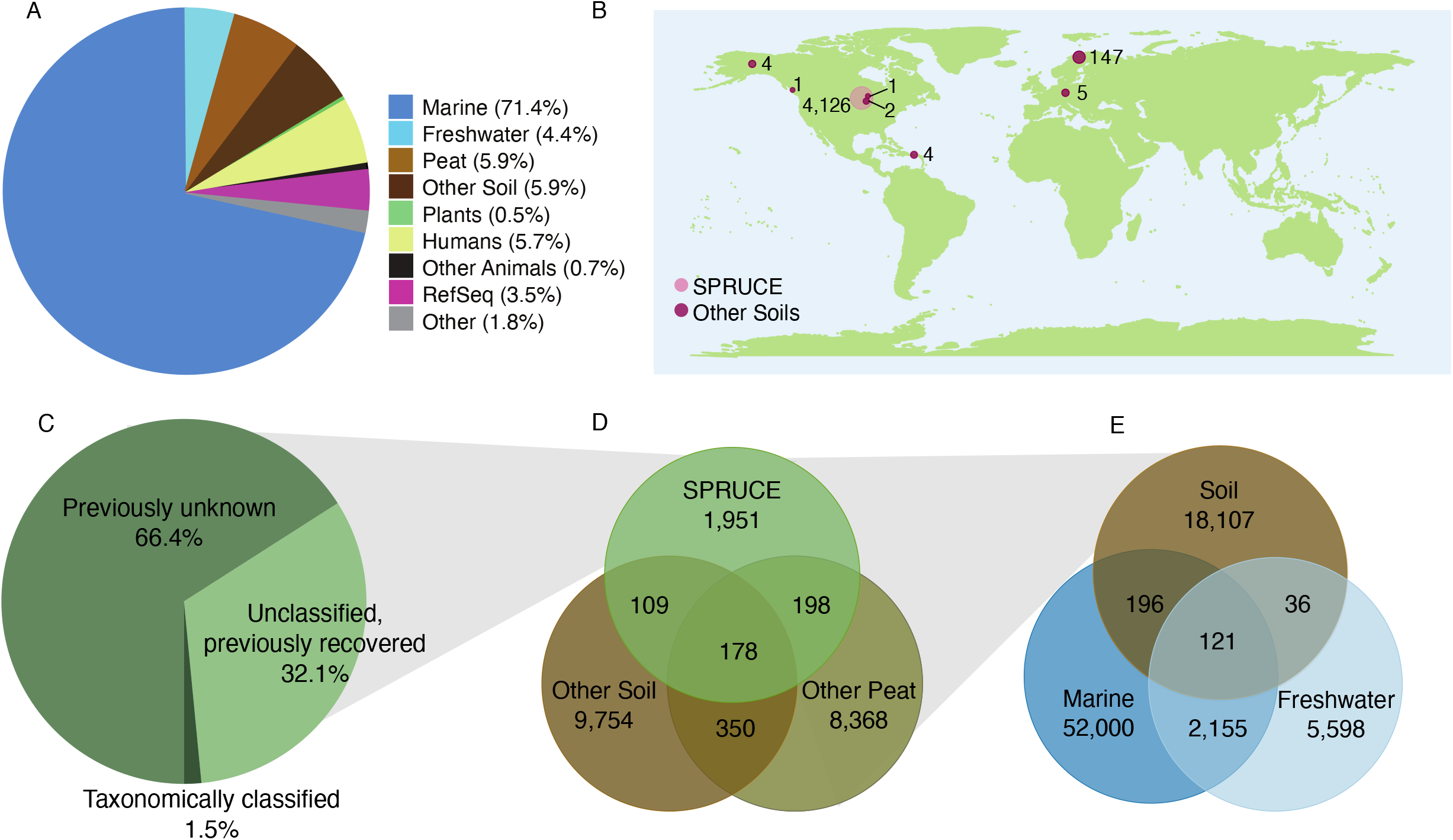
Habitat and global distribution of SPRUCE vOTUs and viral clusters (VCs), using the PIGEON database for context. **A.** Composition of the PIGEON database of vOTUs (n=266,805) by source environment. RefSeq includes isolate viral genomes from a variety of source environments (prokaryotic viruses in RefSeq v95). Plants = plant-associated, Humans = human-associated, Other Animals = non-human animal-associated. **B.** vOTUs (n=4,326) recovered from SPRUCE peat by read mapping, according to the location from which they were first recovered. Numbers indicate SPRUCE vOTUs from a given location. Circle sizes are proportional to the number of vOTUs. **C:** Percentages of vOTUs recovered from SPRUCE that: had predicted taxonomy based on clustering with RefSeq viral genomes (Taxonomically classified), had unknown taxonomy but shared a genus-level viral cluster (VC) with one or more previously recovered vOTUs in the PIGEON database (Unclassified, previously recovered), or were previously unknown at the VC (genus) level (Previously unknown). **D**: Habitat(s) for each soil VC (n=20,908) in the PIGEON database, based on source habitat(s) for the vOTU(s) contained in each VC. For a given soil VC, either all vOTUs were exclusively derived from a single habitat (non-overlapping regions), or two or more vOTUs were derived from different soil habitats (overlapping regions). **E:** Similar to D, but for VCs with vOTUs from soil, marine, and/or freshwater habitats (n=78,213 VCs).

Interestingly, despite the overwhelming dominance of marine vOTUs in our database (190,502 vOTUs, 71%), zero species-level vOTUs from the oceans were recovered in the SPRUCE peatlands. Though freshwater vOTUs (predominantly from freshwater lakes) have less representation in our database (11,869 vOTUs, 4.45%), similarly, no freshwater vOTUs were recovered from SPRUCE peat. Importantly, this analysis at the “species” level is different from the analysis of aquatic-like SPRUCE vOTUs inside the SPRUCE experiment described above; although those were also species-level vOTUs, they were defined (grouped) by shared predicted protein content at the genus level with aquatic vOTUs, such that the same viral “genera” were found in near-surface SPRUCE peatlands and aquatic environments, but none of the SPRUCE vOTUs (“species”) was actually found in aquatic environments. No other vOTUs from our PIGEON database, including bioreactor, hot spring, non-peat wetland, human-, plant-, and other host-associated vOTUs, were recovered in SPRUCE peat. These results suggest viral adaptation to soil and/or strong viral species boundaries between terrestrial, aquatic, and other ecosystems, as previously observed for bacterial species [83,84], though data for soil viruses are limited, so further studies across diverse soils will be necessary to assess the generalizability of these results.

### Taxonomic classification and emergence of global patterns at the “genus” level

To group vOTUs at approximately the genus level, assign taxonomy, and place them in global and ecosystem context, the 4,326 SPRUCE vOTUs were clustered according to shared predicted protein content (using vConTACT2 [85,86]) with the 261,799 other vOTUs in our PIGEON database, including 2,305 RefSeq viral genomes (release 85) [87]. The SPRUCE vOTUs formed 2,445 VCs, 1,457 of which were singletons and 988 of which contained at least two vOTUs (we note that although singletons are not technically clusters, each VC represents a distinct viral “genus” [85,86], so we include singletons in all of our VC counts for ease of interpretation of genus-level trends). Only fourteen of these VCs, containing 67 vOTUs (1.5% of the dataset), were taxonomically classifiable (Fig. 3C), which is substantially less than the taxonomically classifiable portion of previously studied peat viral communities (e.g., 17% of the vOTUs could be taxonomically classified in Emerson et al. 2018 [15]). We speculate that this low level of taxonomic affiliation may be related to the inclusion of more vOTUs from viromes in the current study, relative to the previous work that was focused almost exclusively on viral recovery from total metagenomes. Viromes tend to access more of the rare virosphere [35] and may therefore include vOTUs less likely to be present in the public database used for taxonomic assignments. The taxonomically classifiable vOTUs from SPRUCE included 52 Myoviridae, four Podoviridae, four Siphoviridae, and seven Tectiviridae, consistent with the more abundant viral taxa previously reported from thawing permafrost peatlands [15]. Although most SPRUCE VCs were not taxonomically classifiable, 562 (containing 1,609 vOTUs, 36.6% of the dataset) included a vOTU that was also found in another dataset, meaning that just over 1/3 of the SPRUCE genus-level viral groups had been observed before. The remaining 2,092 SPRUCE VCs (containing 61.8% of the vOTUs) were previously unknown at the genus level.

All 32,346 of the vOTUs from soil in our PIGEON database, including those from SPRUCE and globally distributed soils, grouped into 20,908 genus-level VCs. Of these, 17,488 (83% of the soil VCs, containing 53.9% of the vOTUs) included only a single vOTU, meaning that most of the genus-level viral sequences known from soil worldwide have only been recovered from a single study and/or location so far. In total, 9.3% of the soil VCs, containing 8.2% of the vOTUs, were exclusively found in SPRUCE peatlands. Given that other thoroughly sampled and deeply sequenced peatlands were part of this analysis, these particular viruses may have a limited biogeographical distribution, potentially due to specific adaptations to their local habitats and/or hosts, though further sampling across spatiotemporal scales will be required to more comprehensively unravel local and global peat viral biogeography. Of all of the soil VCs (n=20,908), 178 (0.85%, containing 7.1% of the soil vOTUs) included at least one vOTU each from SPRUCE, other peat habitats, and other soils (Fig. 3D), while 198 VCs (0.94%, 3.1% of the soil vOTUs) contained a vOTU from SPRUCE and other peat sites but not other soils. Together, these data suggest that, while much of soil viral sequence space clearly remains to be explored, genus-level viral similarities may be more common across soil habitats, while species-level similarities may be more restricted to specific soil habitat types.

To investigate similarities between genus-level VCs from soil and aquatic (marine and freshwater) ecosystems, 232,116 vOTUs from our PIGEON database (32,346 soil vOTUs [10,15,31,35], 190,502 vOTUS from marine environments [31,88,89], and 11,869 vOTUs from freshwater environments [31]) were clustered into 78,213 VCs (Table S11). Of the soil VCs, 1.9% shared a cluster with one or both aquatic systems, indicating a small amount of genus-level similarity between aquatic and soil viruses (Fig. 3E). However, most VCs were found in only one habitat, consistent with differences in microbial community composition in aquatic compared to soil and sediment habitats and between freshwater and saltwater environments [83]. Viral clustering according to habitat type has been previously observed, mainly in aquatic viromes, which generally cluster by salinity and other environmental properties [90,91]. Viruses from other ecosystems, such as soil, also tend to be found in similar habitats regardless of geographic location, but this pattern was most pronounced for marine viruses, and comparatively limited data were available from soil [31]. Only 15.4% of the vOTUs from marine environments remained as singleton VCs in our dataset, in contrast with 39.2% of freshwater vOTUs and 45.6% of soil vOTUs. This suggests that marine viral sequence space has been more comprehensively sampled than soil and freshwater habitats, which is not surprising, considering the disproportionate amount of prior research on marine viruses [13,14,17,23,92]. However, repeated sampling of the same kinds of environments (for example, frequent sampling of oxygenated, near-surface photic zones throughout the oceans) would likely yield a similar pattern, even if some habitats (*e.g*., marine oxygen minimum zones) have not been well-sampled. Also, since ocean waters are generally well-mixed and viral populations seem to be transported along ocean currents [45], marine viral populations are presumably more homogeneously dispersed than those in soil or those shared between geographically isolated freshwater bodies [12,18,88,93].

### Comparing viral population (vOTU) recovery from viromes and total soil metagenomes

Metagenomic studies of viral community composition typically take one of two approaches: either the viral signal is mined from total metagenomic assemblies, which predominantly tend to contain bacterial sequencing data [13,15,31], or viral particles are physically separated from other microbes in the laboratory (*e.g*., through filtration), and then viral size-fraction enriched metagenomes (viromes) are sequenced and analyzed [12,13,15,18]. To directly compare results from both approaches, we first analyzed the paired total soil metagenomes and viromes from the five transect samples. Considering all assembled contigs ≥ 10 kbp, only 0.8% of the metagenomic contigs were classified as viral after passing them through viral prediction software (see methods), relative to 16% of the virome contigs. This ~20-fold improvement is consistent with our observed ~30-fold improvement in viral contig recovery from viromes relative to total metagenomes in agricultural soils [35], and similar differences in the composition of metagenomes and viromes have been reported from grassland soils [52]. When accounting for read mapping to all vOTUs in the PIGEON database (including all of the SPRUCE vOTUs), 1,952 vOTUs were detected in the viromes, relative to 401 in the metagenomes from the same samples (Fig. 4A, Supplementary figure 3A). Only 37 vOTUs were detected in the metagenomes alone. Although far more vOTUs were recovered from the viromes, vOTU accumulation curves were still climbing steeply after five samples for both viromes and metagenomes (Fig. 4B, Supplementary figure 3B, 3C), suggesting that more viral diversity remains to be recovered from this peat transect. A comparison of the five viromes indicated that there was no spatial relationship between the samples (Supplementary figure 4A), but there was high variability in the number of recovered vOTUs per sample (Supplementary figure 4B). Notably, sample SPR-2 recovered on average two times more vOTUs than the other viromes, which could be due to a higher sequencing depth, as sample SPR-2 had on average 1.76 times more sequencing than the other viromes.

**Figure 4:**
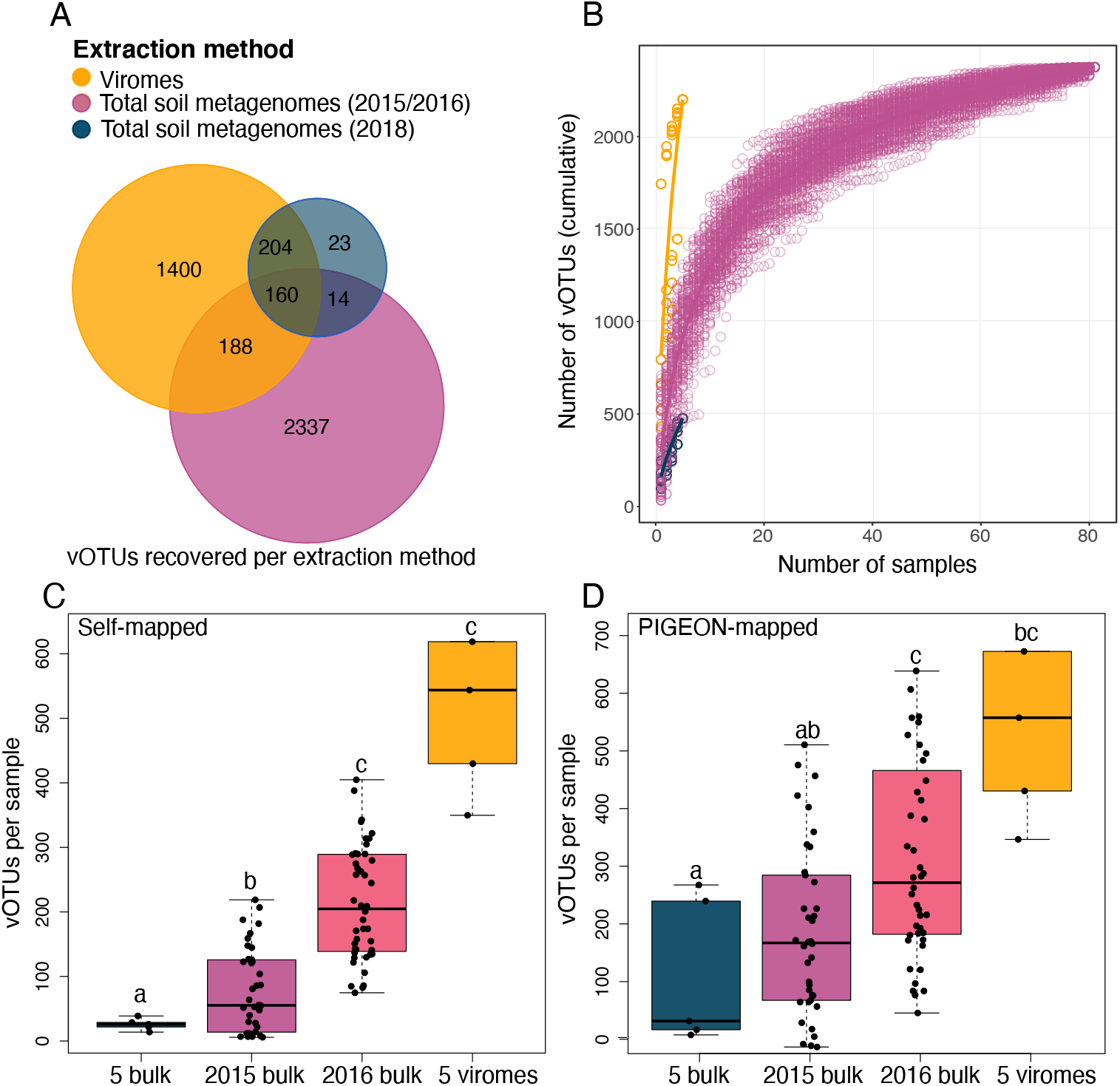
Comparison of vOTU recovery from SPRUCE viromes and total soil metagenomes. **A:** Distribution of vOTUs recovered in each of three extraction groups (grouped by extraction method and collection date), based on read mapping to the PIGEON database (n=5 viromes from 2018, 82 total soil metagenomes from 2015 and 2016, and 5 total soil metagenomes from 2018). **B**: Accumulation curves of distinct vOTUs recovered as sampling increases for each extraction method; 100 permutations of sample order are depicted as open circles, line shows the average of the permutations for each method. **C**: Number of vOTUs recovered per metagenome when reads were only allowed to map to vOTUs that assembled from metagenomes in the same category (self-mapped), considering four categories: 2018 bulk (n=5), 2015 bulk (n=38), 2016 bulk (n=44), 2018 viromes (n=5); bulk = total soil metagenomes. One outlier was excluded from the plot for ease of visualization; the y-axis value of the outlier in the 2018 viromes was 1,328. Letters above boxes correspond to significant differences between groups (Student’s T-test, significant when p < 0.05). **D.** Similar to C, but reads were allowed to map to all vOTUs in the PIGEON database (PIGEON-mapped), including all vOTUs assembled from any of the SPRUCE metagenomes. Three outliers were removed from the plot for ease of visualization; the y-axis values of the two outliers from 2016 bulk were 1,415 and 1,818, and the value of the outlier from the 2018 viromes was 1,558.

To place these direct comparisons of viromes and metagenomes from the same samples in the context of the larger SPRUCE dataset, we compared the five viromes from 2018 to the 82 metagenomes from 2015 and 2016, again with vOTU recovery assessed through read recruitment to all vOTUs in the PIGEON database. We note that the samples in this set of comparisons do differ in multiple ways beyond the extraction method, including the sampling year, depth range, location, and (in some cases) temperature treatment. Specifically, 2015 and 2016 total soil metagenomes were generated from SPRUCE experimental plot samples, most of which received temperature treatments, at four different depths (10-175 cm), whereas the 2018 viromes were recovered from the top 10 cm of a transect outside the experimental plots in the same bog. Also, although all samples were collected in June, the timing of seasonal thaw cycles varies slightly year to year. Acknowledging that all of these sample differences could contribute to the observed trends, on a per-sample basis, the viromes recovered far more vOTUs than the metagenomes, as indicated by the much steeper accumulation curve slope for viromes compared to total metagenomes after only five samples (Fig. 4B). However, the much larger number of samples in the SPRUCE experimental plot metagenomes resulted in a higher total vOTU recovery of 2,699 in the 82 metagenomes, compared to 1,952 in the five viromes (Fig. 4A).

For our final analyses comparing viromes and total metagenomes, we considered the metagenomes from 2015 and 2016 separately, because the sequencing throughput from 2016 was 1.4 times higher than in 2015. The first of these comparisons was based on read recruitment only to vOTUs derived from contigs that assembled from samples in the same category, considering four categories: the five transect viromes, five transect metagenomes, 38 metagenomes from 2015, and 44 metagenomes from 2016. These “self-mapped” analyses were meant to simulate a situation in which only the vOTUs from that particular dataset would have been available. The perceived viral richness per sample was 32 times higher in viromes (mean 649 vOTUs) compared to their five paired metagenomes (mean 20 vOTUs) but was nine and three times higher, respectively, in viromes compared to the 2015 and 2016 metagenomes (mean 72 and 207 vOTUs) (Fig. 4C). The perceived viral richness was 2.8 times higher in the 2016 metagenomes compared to 2015 metagenomes, indicating that a greater sequencing depth of total soil metagenomes (in this case from 6 to 15 Gbp on average) likely increased vOTU recovery, though we cannot exclude the possibility of a true difference in viral richness between the two years. A further comparison of vOTU recovery from the transect viromes and the three sets of metagenomes was based on read recruitment to all 266,805 PIGEON vOTUs from SPRUCE and other datasets. In this case, the perceived viral richness in the viromes (mean 721 vOTUs) was 5.7 times higher than in the paired metagenomes (mean 127 vOTUs, Fig. 4D), 3.5 times higher than in the 2015 metagenomes (mean 200 vOTUs), and two times higher than in the 2016 metagenomes (mean 370 vOTUs). Thus, the availability of reference vOTUs, particularly from the SPRUCE viromes, substantially improved recovery from the total metagenomes.

Few direct comparisons of viromes and total metagenomes from the same samples have been reported from any ecosystem, and even comparisons across different laboratory methods, sequencing throughputs, and numbers of samples are rare. Consistent with our results from peat, agricultural and grassland soil viromes have been shown to be enriched in both viral sequences and genomes from ultrasmall cellular organisms (which would be more likely to pass through the 0.2 μm filters used for viral enrichment) but depleted in sequences from most other cellular organisms, compared to total metagenomes [35,52]. In aqueous systems, water samples are often separated into multiple size fractions (for example, 3-20 μm, 0.8-3 μm, 0.2-0.8 μm, post-0.2 μm), such that previous studies have compared viral sequences recovered across different size fractions, as opposed to comparing the viral fraction to bulk water, and generally, the viruses recovered from different size fractions seem to be distinct [94,95]. A recent meta-analysis of human gut viral data recovered from viromic and metagenomic sequences suggested that more viral contigs could be recovered from metagenomes than from viromes [53]. However, of the 2,017 viromes considered in that study, 1,966 were multiple-displacement amplification (MDA) treated, and, as the authors acknowledged, MDA of viromes has known methodological biases (for example, MDA preferentially recovers circular ssDNA viruses [6]) and thus would result in artificially lower-richness viral communities. Although differences in the environments (human gut compared to soil) could have contributed to the observed differences in viral recovery from viromes compared to total metagenomes in the human gut study compared to our work, the large difference in the number of total metagenomes considered in the human gut study (680) compared to non-MDA amplified viromes (51) could also have contributed to the greater recovery of human gut viral sequences from total metagenomes. Consistent with that interpretation, here we have shown that viromics (without MDA amplification) seems to be a better approach for maximizing viral recovery from soil on a per-sample basis. However, increasing the number of samples, in combination with deeper sequencing and the availability of relevant reference vOTU sequences, improved vOTU recovery from total soil metagenomes, which have the added advantage of accessing virus and host population sequences from the same dataset.

## Conclusions

We analyzed dsDNA viral diversity in a climate-vulnerable peat bog, revealing significant differences in viral community composition at different soil depths and according to peat and porewater C chemistry. Aquatic-like SPRUCE vOTUs were significantly more abundant at near-surface depths, suggesting potential adaptation of these viruses to water-rich environments. Some viral species-level similarities were observed across large geographic distances in soil: 4% of the vOTUs found in SPRUCE peat were previously recovered elsewhere, predominantly in other peatlands, but interestingly, zero marine or freshwater vOTUs were recovered from SPRUCE peat, suggesting the potential for viral species boundaries between terrestrial and aquatic ecosystems. When comparing vOTU recovery from viromes and total soil metagenomes, increasing the dataset size through deeper sequencing and more samples improved vOTU recovery from metagenomes, but viromics was a better approach for maximizing viral recovery on a per-sample basis. Together, these results expand our understanding of soil viral communities and the global soil virosphere, while hinting at a vast diversity of soil viruses remaining to be discovered.

## Materials and methods

### Sample collection

In June 2018, five peat samples were collected along “Transect 4” in the S1 bog ~150 m from the SPRUCE experimental plots in the Marcell Experimental Forest in northern Minnesota, USA (For GPS coordinates, see Table S12). Avoiding green *Sphagnum* moss at the surface (~2 cm), the top 10 cm of peat (5 cm diameter) was collected for each sample with a sterile spatula and placed in 50 mL conical tubes on dry ice. Samples were stored at −80 °C for 6 months prior to DNA extraction for total metagenomes and viromes.

Within the SPRUCE study, temperature treatments were applied in large (~115 sq m) open-topped enclosures. Temperature treatments in the 10 enclosures were as follows: +0, +2.25, +4.5, +6.75 and +9, with two chambers assigned to each temperature treatment. Data were also collected from two ambient environment plots where there was no enclosure but within the treatment area on the south end of the S1 Bog. In each enclosure, warming of deep soil started in June 2014 [55], and aboveground warming began in August 2015 with continuous whole ecosystem warming (365 days per year) operating since late in 2015. A more detailed explanation of deep soil heating procedures and construction of the enclosures and warming mechanics can be found in Hanson et al., 2017 [54,55,62].

Peat samples for 82 total soil metagenomes were collected from the SPRUCE experiment in June 2015 and June 2016 from cores that were extracted using defined hand sampling near the surface and via Russian corers below 30 cm. Samples for analysis were obtained from depth ranges 10-20 cm, 40-50 cm, 100-125 cm, and 150-175 cm from a total of 10 chambers in 2015 (no samples were analyzed from the open, ambient plots that year), with the exception of only two samples collected from chamber 19 (control plot, no temperature treatment, only 10-20 cm and 40-50 cm samples collected), for a total of 38 samples from 2015. In 2016, samples were collected from the same depth ranges from all 10 chambers, plus two samples from each of the two ambient, open plots (depth ranges 10-20 cm and 40-50 cm), for a total of 44 samples from 2016. These 82 samples were used for DNA extraction and total metagenomic analysis and MAG recovery, as described below. Soil temperature, moisture content, CH_4_ and CO_2_ concentrations, and aC measurements (see supplementary methods) were collected from the same samples (Table S13).

### DNA extraction

All samples from the peatland transect were stored at −80°C until further processing. 24 hours prior to DNA extraction, samples were placed at −20 °C. For total metagenomes from the transect, DNA was extracted from 0.25 g peat per sample with the QIAGEN DNeasy Powersoil Kit (QIAGEN, Germany), according to the manufacturer’s protocol. For viromes, 50 g of peat per sample was divided between two 50 mL conical tubes, and 37.5 mL of Amended Potassium Citrate Prime buffer (AKC’, 0.02 μm filtered, 1% K-citrate + 10% PBS + 150 mM MgSO4) [34] was added per tube, for a total of 75 mL buffer. Tubes were shaken at 400 rpm for 15 min, then centrifuged at 4,700 g for 20 min. Excluding the pelleted soil, the supernatant was filtered through a 0.2 μm polyethersulfone filter (Corning, USA) and ultracentrifuged in a Beckman LE-8K ultracentrifuge with a 70 Ti rotor for 3 hours at 32,000 RPM at 4 °C under vacuum. The supernatant was decanted, and the pellet containing virions was resuspended in 200 μl UltraPure water and added to the QIAGEN DNeasy PowerSoil Kit bead tubes (QIAGEN, Germany) for DNA extraction according to the manufacturer’s instructions with one exception: instead of vortexing for 10 minutes with the beads, samples in the bead tubes were incubated at 70 °C for 10 min, vortexed briefly, and incubated at 70 °C for another 5 min. A DNase treatment was not included prior to virion lysis. Anecdotally, this is because we have found that soils stored frozen often have virome DNA yields below detection limits after DNase treatment, while non-DNase-treated viromes from the same frozen samples are still highly virus-enriched relative to total metagenomes (data not shown).

For the 82 2015 and 2016 peat samples used in metagenomic analysis and MAG recovery, DNA was extracted from homogenized samples of each depth interval using the MO BIO Powersoil DNA extraction kit (QIAGEN, Germany). Six replicate 0.35 g extractions were combined and re-purified with the MO BIO PowerClean Pro kit (QIAGEN, Germany) and eluted in 50 mL of 10 mM Tris buffer.

### Library construction and sequencing

Library construction and sequencing for the five viromes and five total soil metagenomes from Transect 4 were conducted by the DNA Technologies and Expression Analysis Cores at the UC Davis Genome Center. Libraries were prepared with the DNA Hyper Prep library kit (Kapa Biosystems-Roche, Basel, Switzerland), as previously described [35]. Paired-end sequencing (150 bp) was done on the Illumina NovaSeq platform, using 4% of a lane per virome and 8% of a lane per total soil metagenome. Sequencing of the 82 metagenomes from the SPRUCE experiment and ambient plots was done by the DOE Joint Genome Institute (JGI), using standard protocols for Nextera XT metagenomic library construction. These barcoded libraries were sequenced on an Illumina HiSeq 2500 instrument in 2×150 bp mode.

### Sequencing read processing, assembly, viral population (vOTU) recovery, and read mapping

Raw reads from the SPRUCE experiment metagenomes (82), transect viromes (5), and transect total soil metagenomes (5) were first quality-trimmed with Trimmomatic v0.38 [96] with a minimum base quality threshold of 30 evaluated on sliding windows of 4 bases and minimum read length of 50. Reads mapped to the PhiX genome were removed with bbduk [97]. Reads were assembled into contigs ≥ 10 kbp in length, using MEGAHIT v 1.1.3 [98] with standard settings. All 92 metagenomes underwent single-sample assemblies, and two additional co-assemblies were generated from the transect, one each for the five viromes and five total soil metagenomes, respectively. For co-assemblies, the preset meta-large option was used. 82 previously existing assemblies from the SPRUCE experiment metagenomes were also used. Briefly, for those assemblies, raw metagenomic fastq sequences were quality trimmed with bbduk from the BBTools software package (options: qtrim=window,2 trimq=17 minlength=100) [99] and assembled with IDBA-UD [100](options: −mink 43 −maxk 123 −step 4 −min_contig 300).

DeepVirFinder [49] and VirSorter [48] were used to recover viral contigs from each assembly. Contigs with DeepVirFinder scores > 0.9 and p < 0.05 were considered viral [88], and DeepVirFinder results were filtered with a custom python script (parse_dvf_results.py, all scripts are available on GitHub, see Data Availability Statement below) to only retain results in compliance with this score. VirSorter was run in regular mode for all total metagenomes and virome decontamination mode for the viromes. Only contigs from VirSorter categories 1, 2, 4 and 5 (high-confidence) were retained. All resulting viral contigs were clustered into vOTUs using CD-HIT [101] at a global identity threshold of 0.95 across 85% of the length of the shorter contig [69]. Different sets of vOTUs were used as references for read mapping throughout the manuscript (see main text), with the most commonly used and most comprehensive reference database being PIGEON (see below). In all cases, read mapping was performed with BBMap [97] at ≥ 90% identity, and vOTU coverage tables were generated with BamM [102], using the ‘tpmean’ setting, and bedfiles were generated using bedtools [103]. Custom python scripts (percentage_coverage.py, filter_coveragetable.py) were used to implement the thresholds for detecting viral populations (vOTUs) in accordance with community standards (≥ 75% of the contig length covered ≥ 1x by reads recruited at ≥ 90% nucleotide identity) [69]. The final vOTU coverage table of per-bp vOTU abundances in each metagenome was normalized by the number of metagenomic sequencing reads for each sample [15].

### Construction of the PIGEON reference database of vOTUs

An in-house database, <u>P</u>hages and <u>I</u>ntegrated <u>G</u>enomes <u>E</u>ncapsidatedOr <u>N</u>ot (PIGEON), was created, containing 266,805 species-level vOTUs, of which 190,502 came from marine environments, 11,869 from freshwater, 32,346 from soil (including 5,006 from SPRUCE), 2,305 RefSeq viral genomes (release 85) [87], and 30,400 from other environments in a meta-analysis, including human microbiomes, other animal microbiomes, plant microbiomes, and other environments). Available viral contigs were downloaded from published datasets [10,13,15,31,34,87–89,104,105], compiled from ongoing work in Alaskan peat soil and Puerto Rican soils (see supplementary methods), and those recovered from SPRUCE (see above). For most of the datasets, viral contigs were derived from viromes, or a combination of viromes and total soil metagenomes, but two datasets only considered viral recovery from total soil metagenomes [10,31]. For all but one of the datasets, VirSorter [48], VirFinder [106], DeepVirFinder [49], or a combination of these programs was used for viral contig recovery (Contigs with DeepVirFinder scores > 0.9 and p < 0.05 were considered viral [88], and only contigs from VirSorter categories 1, 2, 4 and 5 were considered. The exception was the meta-analysis dataset of Paez-Espino et al. (2016), which used a viral discovery pipeline [31]. From all of these datasets, viral contigs ^3^ 10kb were retained and then clustered into vOTUs using CD-HIT [101] at a global identity threshold of 0.95 across 85% of the shorter contig length. PIGEON v1.0 (the version used in this manuscript) is available on Dryad ((https://datadryad.org/, by DOI of this paper). We are actively improving PIGEON and expect to release a new version in the future.

### Viral taxonomic classification and genus-level clustering

Viral taxonomic classifications for the 4,326 SPRUCE vOTUs (detected in the SPRUCE dataset through read mapping) were assigned using vConTACT2 (options: --rel-mode ‘Diamond’ --db ‘ProkaryoticViralRefSeq85-Merged’ −pcs-mode MCL --vcs-mode ClusterONE) [73,74]. The vOTUs were clustered according to shared predicted protein content with the 261,799 other vOTUs in our PIGEON database, including 2,305 RefSeq viral genomes [87]. The viral_cluster_overview output file was used for further analysis, including to manually identify SPRUCE vOTUs that shared a genus-level viral cluster with one or more vOTUs from marine and/or freshwater (aquatic) environments.

### Metagenome-assembled genome (MAG) reconstruction

MAG reconstruction from the five transect total metagenomes was done as follows: quality-trimmed reads were assembled using MEGAHITv 1.1.3 [98] with a minimum contig length of 2,000, using the meta-large preset. After individual assembly of each sample, quality-filtered and trimmed reads were mapped to the resulting contigs using bbmap [107] with standard settings, and this abundance information was used to bin the contigs into MAGs using MetaBAT [108], using the --veryspecific setting and the coverage depth information. Quality and identification of bins was done with CheckM [109], following Sorensen et al., [110].

From the 82 SPRUCE experiment metagenomes, metagenome assembly, recovery, and analysis of metagenome-assembled genomes (MAGs) was performed as described in Johnston et al., [111]. Briefly, metagenomic sequences were assembled with IDBA-UD [100] (options: - mink 43 −maxk 123 −step 4 −min_contig 300). Resulting contigs ≥ 2.5 kbp were used to recover microbial population genomes with MetaBAT2 (options: –minCVSum 10) [108] and MaxBin2 [112]. Before binning, Bowtie 2 was used to align short-read sequences to assembled contigs (options: −very-fast) [113], and SAMtools was used to sort and convert SAM files to BAM format [114]. Sorted BAM files were then used to calculate the coverage (mean representation) of each contig in each metagenome. The quality of each resulting MAG was evaluated with the CheckM v1.0.3 taxonomy workflow for Bacteria and Archaea separately [109]. The result from either evaluation (i.e., taxonomy workflow for Archaea or Bacteria) with the highest estimated completeness was retained for each MAG. MAGs with a quality score ≥ 60 were retained (from Parks et al., 2017 [115] calculated as the estimated completeness – 5 × contamination). MAGs recovered from different metagenomes were dereplicated with dREP [116], and the GTDB-tk classify workflow [117,118] was used to determine MAG taxonomic affiliations. MAG gene prediction, functional annotation, and assessment of metabolic pathway completeness (e.g., for assessing methanogenesis potential) was performed as described in Johnston et al., 2019 [111]. Taxonomic classification, source dataset SRA ID, basic genome statistics, and CheckM summaries for each MAG can be found in Table S4.

Using the parameters described above for vOTU coverage table generation, a microbial contig coverage table was generated. From this coverage table, we calculated the coverage of each population genome as the average of all of its binned contig coverages, weighting each contig by its length in base pairs. In-house scripts for this are available on GitHub. Hmm searches were done on both MAGs and vOTUs for proteins involved in methanogenesis or methanotrophy (McrA (a methanogenesis biomarker) [75,76], sMMO, pMMO, and pXMO (methanotrophy biomarkers) [3]). The MAG and vOTU contigs were annotated with prodigal (standard settings) [119], and an HMM search was done on these annotations with hmmr [120], using hmmsearch (standard settings) with an e-value cutoff of 1E-5 [121].

### Reconstruction of microbial CRISPR arrays and virus-host linkages

CRISPR repeat and spacer arrays were assembled with Crass v0.3.12 [74], using standard settings, and BLASTn was used to match spacer sequences with vOTUs and repeats to MAGs, in order to link viruses to putative hosts. Briefly, for protospacer-spacer matches (*i.e*., matches between vOTUs and CRISPR spacer sequences), the BLASTn-short function was used, with £ 1 mismatch to spacer sequences, e-value threshold of 1.0×10^-10^, and a percent identity of 95 [31,122]. For MAG-repeat matches, the BLASTn-short function was used, with an e-value threshold of 1.0×10^-10^ and a percent identity of 100 [15].

### Phylogenetic tree construction

A phylogenetic tree of bacterial host MAGs with CRISPR matches to one or more vOTUs (*i.e*., a repeat match to a MAG and a spacer from the same CRISPR array with a match to a vOTU protospacer) was constructed with CheckM [109] via a marker-gene alignment of 43 conserved marker genes with largely congruent phylogenetic histories, defined by CheckM [109]. This alignment was used to construct a maximum-likelihood tree with MEGA [123], with the LG plus frequencies model [124]. A total of 500 bootstrap replicates were conducted under the neighbor-joining method with a Poisson model.

### Data analysis (ecological statistics)

The following statistical analyses were performed in R using the Vegan [125] package: accumulation curves were calculated using the speccacum function, vOTU coverage tables were standardized using the decostand function with the Hellinger method, and Bray-Curtis dissimilarity matrices were calculated using the vegdist function. Mantel tests were performed with the mantel function, using the Pearson method, and permutational multivariate analyses of variance (PERMANOVA) were performed with the Adonis function. Venn diagrams were created with the VennDiagram package, using the draw.triple.venn function. The differential abundance analysis of vOTUs across depth levels was performed using the likelihood ratio test implemented in DESeq2 [126]. Hierarchical clustering of the viral abundance patterns of the five viromes was done with the hclust function (method=complete), and heatmaps were created with the pheatmap and dendextend libraries. The world map was created with the maps library.

### Detection of putative viral auxiliary metabolic genes (AMGs)

VIBRANT [50] and DRAM-v [51] were used to identify putative AMGs in the vOTU sequences. VIBRANT was run (using standard settings) on all SPRUCE viral contigs identified by either VirSorter or DeepVirFinder, resulting in 2,802 vOTUs that were used for this analysis. VIBRANT output was manually screened to determine whether the predicted AMGs had viral genes upstream and downstream [15], and in many cases, they did not (see supplementary discussion). DRAM-v (standard settings) was applied to 2,645 vOTUs that were recovered by both VIBRANT and VirSorter, because DRAM-v uses the VirSorter output, and we wanted to compare results from the two AMG detection methods. From the DRAM-v output, only putative AMGs with auxiliary scores < 4 were retained (a low auxiliary score indicates a gene that is confidently viral), and no viral flag (F), transposon flag (T), viral-like peptidase (P), or attachment flag (A) could be present. Putative AMGs that did not have a gene ID or a gene description were also discarded. See supplemental discussion for more information.

## Supporting information

supplemental materals, methods, discussion and figures

## Declarations

### Ethics approval and consent to participate

Not applicable

### Consent for publication

Not applicable

### Availability of data and material

The raw sequencing datasets from the SPRUCE transect have been deposited in the Sequence Read Archive (BioProject PRJNA666221). The 5,006 vOTUs from SPRUCE, the 486 MAGs from SPRUCE and the PIGEON database are available at Dryad (https://datadryad.org/, by DOI of this paper). Sequencing data from the 82 SPRUCE experiment metagenomes were downloaded from the SPRUCE website (https://mnspruce.ornl.gov/node/622, https://mnspruce.ornl.gov/node/727, accessed June 2019, Table S13), where they are currently still available. In addition, these 82 metagenomes are available from the JGI Genome Portal and NCBI Sequence Read Archive (SRA). SRA identifiers for each metagenomic dataset are provided in Table S13. Relevant processed data and geochemical data are available as Tables S12 and S13. Code for processing viromic data and all relevant R and python scripts are available on GitHub (https://github.com/AnneliektH/SPRUCE)

### Authors’ contributions

AMH and JBE designed the study and wrote the manuscript. JBE collected and AMH processed the 2018 transect samples. RMW generated geochemical data. AMH, CSM, JWS, LAZ, RMW, ERJ, and JBE performed data analysis. GGT, SJB, and JPR contributed vOTU sequences to the PIGEON database from their ongoing work in Alaskan and Puerto Rican soils. RMW, PJH, JPC, CWS, and JEK facilitated field site and/or data access and integration and were liaisons to the larger SPRUCE project. All authors contributed to project discussions, edited the manuscript, and approved the final version of the manuscript.

## Acknowledgements

We thank Sara Geonczy for helpful comments on the manuscript, Winston Bess and Rose Bolle for assistance in preparing for field work, and Sarah Lutman, Robert Rudolph, and Margaret Rudolph for handling shipments and logistical support en route to the field. We thank Alena Schroeder and Gerdie ter Horst for helpful contributions to project discussions. For the Puerto Rico viral sequences, we thank Ashley Campbell, Amrita Bhattacharyya, and Jeff Kimbrel for carrying out the original experiment and processing the metagenomes.

## Notice

Effort contributing to this manuscript has been authored in part by UT-Battelle, LLC under Contract No. DE-AC05-00OR22725 with the US Department of Energy. The United States Government retains and the publisher, by accepting the article for publication, acknowledges that the United States Government retains a non-exclusive, paid-up, irrevocable, world-wide license to publish or reproduce the published form of this manuscript, or allow others to do so, for United States Government purposes. The Department of Energy will provide public access to these results of federally sponsored research in accordance with the DOE Public Access Plan (http://energy.gov/downloads/doe-public-access-plan).

## Supplementary Figures

**Supplementary figure 1: Sampling locations for all SPRUCE samples.** Sampling locations within the S1 Bog at the Marcell Experimental Forest in Northern Minnesota, USA, including the five transect samples and the samples from the SPRUCE experimental plots and chambers. Numbers next to the brackets show how many and what kinds of metagenomes were derived from each part of the bog.

**Supplementary figure 2: Comparison of vOTU recovery from five paired viromes and total soil metagenomes from the SPRUCE transect. A:** Distribution of vOTUs recovered by each of the two extraction methods, based on read mapping to the PIGEON database, including all vOTUs recovered from SPRUCE. **B:** Accumulation curves of distinct vOTUs recovered as sampling increases for each extraction method; 100 permutations of sample order are depicted as open circles, and averages are shown as a line. **C:** Similar to panel B, but only the accumulation curve of distinct vOTUs recovered from total soil metagenomes is shown, with a smaller y-axis maximum to better show the trend.

**Supplementary figure 3: Comparison of the five viromes from the transect. A:** Dendrogram depicting sample similarity according to viral community composition (left) and heatmap (right) of vOTUs detected (green = detected, white = not detected) in the five SPRUCE transect viromes. **B**: Comparison of vOTU recovery from the SPRUCE-2 sample compared to the four other virome samples.

## Additional files

Additional file 1

Figure_S1.png

Figure S1

Sampling locations for all SPRUCE samples

Additional file 2

Figure_S2.pdf

Figure S2

Comparison of vOTU recovery from five paired viromes and total soil metagenomes from the SPRUCE transect

Additional file 3

Figure_S3.pdf

Figure S3

Comparison of the five viromes from the transect

Additional file 4

SPRUCE_supplemental_tables.xlsx

Excel file (xlxs)

Tables S1-S14

All supplemental tables that are referenced in the text. Each sheet is a separate supplemental table.

Additional file 5

201214_SPRUCE_supplementaldata.docx

Word document (docx)

Supplemental discussion and methods

Supplemental discussion and methods text for this manuscript.

## References

1. Wilson RM, Hopple AM, Tfaily MM, Sebestyen SD, Schadt CW, Pfeifer-Meister L, et al. Stability of peatland carbon to rising temperatures. Nat Commun. 2016;7:13723.

2. Tveit AT, Urich T, Svenning MM. Metatranscriptomic analysis of arctic peat soil microbiota. Appl Environ Microbiol. 2014;80:5761–72.

3. Singleton CM, McCalley CK, Woodcroft BJ, Boyd JA, Evans PN, Hodgkins SB, et al. Methanotrophy across a natural permafrost thaw environment. ISME J. 2018;12:2544–58.

4. Mondav R, Woodcroft BJ, Kim E-H, McCalley CK, Hodgkins SB, Crill PM, et al. Discovery of a novel methanogen prevalent in thawing permafrost. Nature Communications. 2014. Available from: http://dx.doi.org/10.1038/ncomms4212

5. Schuur EAG, McGuire AD, Schädel C, Grosse G, Harden JW, Hayes DJ, et al. Climate change and the permafrost carbon feedback. Nature. 2015;520:171–9.

6. Williamson KE, Fuhrmann JJ, Wommack KE, Radosevich M. Viruses in Soil Ecosystems: An Unknown Quantity Within an Unexplored Territory. Annu Rev Virol. 2017;4:201–19.

7. Trubl G, Solonenko N, Chittick L, Solonenko SA, Rich VI, Sullivan MB. Optimization of viral resuspension methods for carbon-rich soils along a permafrost thaw gradient. PeerJ. 2016;4:e1999.

8. Narr A, Nawaz A, Wick LY, Harms H, Chatzinotas A. Soil viral communities vary temporally and along a land use transect as revealed by virus-like particle counting and a modified community fingerprinting approach (fRAPD). Front Microbiol. Frontiers; 2017;8:1975.

9. Williamson KE, Corzo KA, Drissi CL, Buckingham JM, Thompson CP, Helton RR. Estimates of viral abundance in soils are strongly influenced by extraction and enumeration methods. Biol Fertil Soils. Springer; 2013;49:857–69.

10. Dalcin Martins P, Danczak RE, Roux S, Frank J, Borton MA, Wolfe RA, et al. Viral and metabolic controls on high rates of microbial sulfur and carbon cycling in wetland ecosystems. Microbiome. 2018;6:138.

11. Hurwitz BL, U’Ren JM. Viral metabolic reprogramming in marine ecosystems. Curr Opin Microbiol. 2016;31:161–8.

12. Roux S, Brum JR, Dutilh BE, Sunagawa S, Duhaime MB, Loy A, et al. Ecogenomics and potential biogeochemical impacts of globally abundant ocean viruses. Nature. 2016;537:689–93.

13. Trubl G, Jang HB, Roux S, Emerson JB, Solonenko N, Vik DR, et al. Soil Viruses Are Underexplored Players in Ecosystem Carbon Processing. mSystems. 2018. Available from: http://dx.doi.org/10.1128/msystems.00076-18

14. Emerson JB. Soil Viruses: A New Hope. mSystems. 2019;4. Available from: http://dx.doi.org/10.1128/mSystems.00120-19

15. Emerson JB, Roux S, Brum JR, Bolduc B, Woodcroft BJ, Jang HB, et al. Host-linked soil viral ecology along a permafrost thaw gradient. Nature Microbiology. Nature Publishing Group; 2018;3:870–80.

16. Sieradzki ET, Ignacio-Espinoza JC, Needham DM, Fichot EB, Fuhrman JA. Dynamic marine viral infections and major contribution to photosynthetic processes shown by spatiotemporal picoplankton metatranscriptomes. Nat Commun. 2019;10:1169.

17. Breitbart M, Bonnain C, Malki K, Sawaya NA. Phage puppet masters of the marine microbial realm. Nat Microbiol. 2018;3:754–66.

18. Brum JR, Sullivan MB. Rising to the challenge: accelerated pace of discovery transforms marine virology. Nat Rev Microbiol. 2015;13:147–59.

19. Fierer N. Embracing the unknown: disentangling the complexities of the soil microbiome. Nat Rev Microbiol. 2017;15:579–90.

20. Pratama AA, van Elsas JD. The “Neglected” Soil Virome - Potential Role and Impact. Trends Microbiol. 2018;26:649–62.

21. Kuzyakov Y, Mason-Jones K. Viruses in soil: Nano-scale undead drivers of microbial life, biogeochemical turnover and ecosystem functions. Soil Biol Biochem. 2018;127:305–17.

22. Williamson KE, Wommack KE, Radosevich M. Sampling natural viral communities from soil for culture-independent analyses. Appl Environ Microbiol. 2003;69:6628–33.

23. Williamson KE, Radosevich M, Wommack KE. Abundance and diversity of viruses in six Delaware soils. Appl Environ Microbiol. 2005;71:3119–25.

24. Williamson KE, Radosevich M, Smith DW, Wommack KE. Incidence of lysogeny within temperate and extreme soil environments. Environ Microbiol. 2007;9:2563–74.

25. Swanson MM, Fraser G, Daniell TJ, Torrance L, Gregory PJ, Taliansky M. Viruses in soils: morphological diversity and abundance in the rhizosphere. Ann Appl Biol. 2009;155:51–60.

26. Ghosh D, Roy K, Williamson KE, Srinivasiah S, Wommack KE, Radosevich M. Acylhomoserine lactones can induce virus production in lysogenic bacteria: an alternative paradigm for prophage induction. Appl Environ Microbiol. 2009;75:7142–52.

27. Liu J, Yu Z, Wang X, Jin J, Liu X, Wang G. The distribution characteristics of the major capsid gene (g23) of T4-type phages in paddy floodwater in Northeast China. Soil Sci Plant Nutr. Taylor & Francis; 2016;62:133–9.

28. Zablocki O, van Zyl L, Adriaenssens EM, Rubagotti E, Tuffin M, Cary SC, et al. High-level diversity of tailed phages, eukaryote-associated viruses, and virophage-like elements in the metaviromes of antarctic soils. Appl Environ Microbiol. 2014;80:6888–97.

29. Williamson KE, Schnitker JB, Radosevich M, Smith DW, Wommack KE. Cultivation-based assessment of lysogeny among soil bacteria. Microb Ecol. 2008;56:437–47.

30. Ghosh D, Roy K, Williamson KE, White DC, Wommack KE, Sublette KL, et al. Prevalence of lysogeny among soil bacteria and presence of 16S rRNA and trzN genes in viral-community DNA. Appl Environ Microbiol. 2008;74:495–502.

31. Paez-Espino D, Eloe-Fadrosh EA, Pavlopoulos GA, Thomas AD, Huntemann M, Mikhailova N, et al. Uncovering Earth’s virome. Nature. 2016;536:425–30.

32. Starr EP, Nuccio EE, Pett-Ridge J, Banfield JF, Firestone MK. Metatranscriptomic reconstruction reveals RNA viruses with the potential to shape carbon cycling in soil. Proc Natl Acad Sci U S A. 2019;116:25900–8.

33. Stough JMA, Kolton M, Kostka JE, Weston DJ, Pelletier DA, Wilhelm SW. Diversity of Active Viral Infections within the Sphagnum Microbiome. Appl Environ Microbiol. 2018;84. Available from: http://dx.doi.org/10.1128/AEM.01124-18

34. Trubl G, Roux S, Solonenko N, Li Y-F, Bolduc B, Rodríguez-Ramos J, et al. Towards optimized viral metagenomes for double-stranded and single-stranded DNA viruses from challenging soils. PeerJ. 2019;7:e7265.

35. Santos-Medellin C, Zinke LA, ter Horst AM, Gelardi DL, Parikh SJ, Emerson JB. Viromes outperform total metagenomes in revealing the spatiotemporal patterns of agricultural soil viral communities. 2020. p. 2020.08.06.237214. Available from: https://www.biorxiv.org/content/10.1101/2020.08.06.237214v1.abstract

36. Mackelprang R, Saleska SR, Jacobsen CS, Jansson JK, Taş N. Permafrost Meta-Omics and Climate Change. Annu Rev Earth Planet Sci. Annual Reviews; 2016;44:439–62.

37. Jansson JK, Taş N. The microbial ecology of permafrost. Nat Rev Microbiol. 2014;12:414–25.

38. Woodcroft BJ, Singleton CM, Boyd JA, Evans PN, Emerson JB, Zayed AAF, et al. Genome-centric view of carbon processing in thawing permafrost. Nature. 2018;560:49–54.

39. Lin X, Tfaily MM, Steinweg JM, Chanton P, Esson K, Yang ZK, et al. Microbial community stratification linked to utilization of carbohydrates and phosphorus limitation in a boreal peatland at Marcell Experimental Forest, Minnesota, USA. Appl Environ Microbiol. 2014;80:3518–30.

40. Lin X, Tfaily MM, Green SJ, Steinweg JM, Chanton P, Imvittaya A, et al. Microbial metabolic potential for carbon degradation and nutrient (nitrogen and phosphorus) acquisition in an ombrotrophic peatland. Appl Environ Microbiol. 2014;80:3531–40.

41. Tfaily MM, Cooper WT, Kostka JE, Chanton PR, Schadt CW, Hanson PJ, et al. Organic matter transformation in the peat column at Marcell Experimental Forest: Humification and vertical stratification. Journal of Geophysical Research: Biogeosciences. John Wiley & Sons, Ltd; 2014;119:661–75.

42. Tfaily MM, Wilson RM, Cooper WT, Kostka JE, Hanson P, Chanton JP. Vertical Stratification of Peat Pore Water Dissolved Organic Matter Composition in a Peat Bog in Northern Minnesota. Journal of Geophysical Research: Biogeosciences. John Wiley & Sons, Ltd; 2018;123:479–94.

43. Mackelprang R, Waldrop MP, DeAngelis KM, David MM, Chavarria KL, Blazewicz SJ, et al. Metagenomic analysis of a permafrost microbial community reveals a rapid response to thaw. Nature. 2011;480:368–71.

44. Hultman J, Waldrop MP, Mackelprang R, David MM, McFarland J, Blazewicz SJ, et al. Multi-omics of permafrost, active layer and thermokarst bog soil microbiomes. Nature. 2015;521:208–12.

45. Brum JR, Ignacio-Espinoza JC, Roux S, Doulcier G, Acinas SG, Alberti A, et al. Ocean plankton. Patterns and ecological drivers of ocean viral communities. Science. 2015;348:1261498.

46. Göller PC, Haro-Moreno JM, Rodriguez-Valera F, Loessner MJ, Gómez-Sanz E. Uncovering a hidden diversity: optimized protocols for the extraction of dsDNA bacteriophages from soil. Microbiome. 2020. Available from: http://dx.doi.org/10.1186/s40168-020-0795-2

47. Trubl G, Hyman P, Roux S, Abedon ST. Coming-of-Age Characterization of Soil Viruses: A User’s Guide to Virus Isolation, Detection within Metagenomes, and Viromics. Soil Systems. 2020. p. 23. Available from: http://dx.doi.org/10.3390/soilsystems4020023

48. Roux S, Enault F, Hurwitz BL, Sullivan MB. VirSorter: mining viral signal from microbial genomic data. PeerJ. 2015;3:e985.

49. Ren J, Song K, Deng C, Ahlgren NA, Fuhrman JA, Li Y, et al. Identifying viruses from metagenomic data using deep learning. Quantitative Biology. 2020;8:64–77.

50. Kieft K, Zhou Z, Anantharaman K. VIBRANT: automated recovery, annotation and curation of microbial viruses, and evaluation of viral community function from genomic sequences. Microbiome. 2020;8:90.

51. Shaffer M, Borton MA, McGivern BB, Zayed AA, La Rosa SL, Solden LM, et al. DRAM for distilling microbial metabolism to automate the curation of microbiome function. Nucleic Acids Res. 2020; Available from: http://dx.doi.org/10.1093/nar/gkaa621

52. Nicolas AM, Jaffe AL, Nuccio EE, Taga ME, Firestone MK, Banfield JF. Unexpected diversity of CPR bacteria and nanoarchaea in the rare biosphere of rhizosphere-associated grassland soil. Cold Spring Harbor Laboratory. 2020. p. 2020.07.13.194282. Available from: https://www.biorxiv.org/content/10.1101/2020.07.13.194282v1

53. Gregory AC, Zablocki O, Zayed AA, Howell A, Bolduc B, Sullivan MB. The Gut Virome Database Reveals Age-Dependent Patterns of Virome Diversity in the Human Gut. Cell Host Microbe. 2020; Available from: http://dx.doi.org/10.1016/j.chom.2020.08.003

54. Norby RJ, Childs J, Hanson PJ, Warren JM. Rapid loss of an ecosystem engineer: Sphagnum decline in an experimentally warmed bog. Ecol Evol. 2019;9:12571–85.

55. Hanson PJ, Riggs JS, Nettles WR, Phillips JR, Krassovski MB, Hook LA, et al. Attaining whole-ecosystem warming using air and deep-soil heating methods with an elevated CO_2_ atmosphere. Biogeosciences. Copernicus GmbH; 2017;14:861–83.

56. Dise NB, Gorham E, Verry ES. Environmental factors controlling methane emissions from peatlands in northern Minnesota. J Geophys Res. 1993;98:10583.

57. Kolka R, Sebestyen S, Verry ES, Brooks K. Peatland Biogeochemistry and Watershed Hydrology at the Marcell Experimental Forest. CRC Press; 2011.

58. Grigal DF. Elemental dynamics in forested bogs in northern Minnesota. Can J Bot. NRC Research Press; 1991;69:539–46.

59. Nichols DS, Brown JM. Evaporation from a sphagnum moss surface. J Hydrol. 1980;48:289–302.

60. Verry ES, Timmons DR. Waterborne Nutrient Flow Through an Upland-Peatland Watershed in Minnesota. Ecology. 1982. p. 1456–67. Available from: http://dx.doi.org/10.2307/1938872

61. Boelter DH, Verry ES. Peatland and Water in the Northern Lake States. Department of Agriculture, Forest Service, North Central Forest Experiment Station; 1977.

62. Richardson AD, Hufkens K, Milliman T, Aubrecht DM, Furze ME, Seyednasrollah B, et al. Ecosystem warming extends vegetation activity but heightens vulnerability to cold temperatures. Nature. 2018;560:368–71.

63. Fernandez CW, Heckman K, Kolka R, Kennedy PG. Melanin mitigates the accelerated decay of mycorrhizal necromass with peatland warming. Ecol Lett. 2019;22:498–505.

64. McPartland MY, Kane ES, Falkowski MJ, Kolka R, Turetsky MR, Palik B, et al. The response of boreal peatland community composition and NDVI to hydrologic change, warming, and elevated carbon dioxide. Glob Chang Biol. 2019;25:93–107.

65. Hopple AM, Wilson RM, Kolton M, Zalman CA, Chanton JP, Kostka J, et al. Massive peatland carbon banks vulnerable to rising temperatures. Nat Commun. 2020;11:2373.

66. Carrell AA, Kolton M, Glass JB, Pelletier DA, Warren MJ, Kostka JE, et al. Experimental warming alters the community composition, diversity, and N2 fixation activity of peat moss (Sphagnum fallax) microbiomes. Glob Chang Biol. 2019;25:2993–3004.

67. Warren MJ, Lin X, Gaby JC, Kretz CB, Kolton M, Morton PL, et al. Molybdenum-Based Diazotrophy in a Sphagnum Peatland in Northern Minnesota. Applied and Environmental Microbiology. 2017. Available from: http://dx.doi.org/10.1128/aem.01174-17

68. Kluber LA, Johnston ER, Allen SA, Hendershot JN, Hanson PJ, Schadt CW. Constraints on microbial communities, decomposition and methane production in deep peat deposits. PLoS One. 2020;15:e0223744.

69. Roux S, Adriaenssens EM, Dutilh BE, Koonin EV, Kropinski AM, Krupovic M, et al. Minimum Information about an Uncultivated Virus Genome (MIUViG). Nat Biotechnol. 2019;37:29–37.

70. Liang X, Wagner RE, Zhuang J, DeBruyn JM, Wilhelm SW, Liu F, et al. Viral abundance and diversity vary with depth in a southeastern United States agricultural ultisol. Soil Biol Biochem. 2019;137:107546.

71. McCalley CK, Woodcroft BJ, Hodgkins SB, Wehr RA, Kim E-H, Mondav R, et al. Methane dynamics regulated by microbial community response to permafrost thaw. Nature. 2014;514:478–81.

72. Hodgkins SB, Chanton JP, Langford LC, McCalley CK, Saleska SR, Rich VI, et al. Soil incubations reproduce field methane dynamics in a subarctic wetland. Biogeochemistry. 2015;126:241–9.

73. Hobbie EA, Chen J, Hanson PJ, Iversen CM, McFarlane KJ, Thorp NR, et al. Long-term carbon and nitrogen dynamics at SPRUCE revealed through stable isotopes in peat profiles. Biogeosciences. 2017. p. 2481–94. Available from: http://dx.doi.org/10.5194/bg-14-2481-2017

74. Skennerton CT, Imelfort M, Tyson GW. Crass: identification and reconstruction of CRISPR from unassembled metagenomic data. Nucleic Acids Res. 2013;41:e105.

75. Evans PN, Boyd JA, Leu AO, Woodcroft BJ, Parks DH, Hugenholtz P, et al. An evolving view of methane metabolism in the Archaea. Nat Rev Microbiol. 2019;17:219–32.

76. Zinke LA, Evans PN, Schroeder AL, Parks DH, Varner RK, Rich VI, et al. 1 Evidence for non-methanogenic metabolisms in globally distributed archaeal clades basal 2 to the Methanomassiliicoccales. Available from: http://dx.doi.org/10.1101/2020.03.09.984617

77. Jin M, Guo X, Zhang R, Qu W, Gao B, Zeng R. Diversities and potential biogeochemical impacts of mangrove soil viruses. Microbiome. 2019;7:58.

78. Du Toit A. Permafrost thawing and carbon metabolism. Nat. Rev. Microbiol. 2018. p. 519.

79. DeAngelis KM, Pold G, Topçuoğlu BD, van Diepen LTA, Varney RM, Blanchard JL, et al. Long-term forest soil warming alters microbial communities in temperate forest soils. Front Microbiol. 2015;6:104.

80. Liu D, Keiblinger KM, Schindlbacher A, Wegner U, Sun H, Fuchs S, et al. Microbial functionality as affected by experimental warming of a temperate mountain forest soil—A metaproteomics survey. Appl Soil Ecol. 2017;117-118:196–202.

81. Wang H, Liu S, Schindlbacher A, Wang J, Yang Y, Song Z, et al. Experimental warming reduced topsoil carbon content and increased soil bacterial diversity in a subtropical planted forest. Soil Biol Biochem. 2019;133:155–64.

82. Kolton M, Marks A, Wilson RM, Chanton JP, Kostka JE. Impact of Warming on Greenhouse Gas Production and Microbial Diversity in Anoxic Peat From a Sphagnum-Dominated Bog (Grand Rapids, Minnesota, United States). Front Microbiol. 2019;10:870.

83. Lozupone CA, Knight R. Global patterns in bacterial diversity. Proc Natl Acad Sci U S A. 2007;104:11436–40.

84. Thompson LR, Sanders JG, McDonald D, Amir A, Ladau J, Locey KJ, et al. A communal catalogue reveals Earth’s multiscale microbial diversity. Nature. 2017;551:457–63.

85. Bolduc B, Jang HB, Doulcier G, You Z-Q, Roux S, Sullivan MB. vConTACT: an iVirus tool to classify double-stranded DNA viruses that infect Archaea and Bacteria. PeerJ. 2017;5:e3243.

86. Bin Jang H, Bolduc B, Zablocki O, Kuhn JH, Roux S, Adriaenssens EM, et al. Taxonomic assignment of uncultivated prokaryotic virus genomes is enabled by gene-sharing networks. Nat Biotechnol. 2019;37:632–9.

87. Pruitt KD, Tatusova T, Maglott DR. NCBI reference sequences (RefSeq): a curated non-redundant sequence database of genomes, transcripts and proteins. Nucleic Acids Res. 2007;35:D61–5.

88. Gregory AC, Zayed AA, Conceição-Neto N, Temperton B, Bolduc B, Alberti A, et al. Marine DNA Viral Macro- and Microdiversity from Pole to Pole. Cell. 2019;177:1109–23.e14.

89. Roux S, Hallam SJ, Woyke T, Sullivan MB. Viral dark matter and virus–host interactions resolved from publicly available microbial genomes. Elife. eLife Sciences Publications, Ltd; 2015;4:e08490.

90. Roux S, Enault F, Ravet V, Colombet J, Bettarel Y, Auguet J-C, et al. Analysis of metagenomic data reveals common features of halophilic viral communities across continents. Environ Microbiol. 2016;18:889–903.

91. Emerson JB. Assembly of Deeply Sequenced Metagenomes Yields Insight into Viral and Microbial Ecology in Two Natural Systems. UC Berkeley; 2012. Available from: https://escholarship.org/uc/item/321735jt

92. Breitbart M, Thompson LR, Suttle CA, Sullivan MB. Exploring the Vast Diversity of Marine Viruses. Oceanography. Oceanography Society; 2007;20:135–9.

93. Chow C-ET, Suttle CA. Biogeography of Viruses in the Sea. Annu Rev Virol. 2015;2:41–66.

94. Williamson SJ, Allen LZ, Lorenzi HA, Fadrosh DW, Brami D, Thiagarajan M, et al. Metagenomic exploration of viruses throughout the Indian Ocean. PLoS One. 2012;7:e42047.

95. Emerson JB, Andrade K, Thomas BC, Norman A, Allen EE, Heidelberg KB, et al. Virus-host and CRISPR dynamics in Archaea-dominated hypersaline Lake Tyrrell, Victoria, Australia. Archaea. 2013;2013:370871.

96. Bolger AM, Lohse M, Usadel B. Trimmomatic: a flexible trimmer for Illumina sequence data. Bioinformatics. 2014;30:2114–20.

97. Bushnell B. BBTools software package. URL http://sourceforgenet/projects/bbmap. 2014;

98. Li D, Liu C-M, Luo R, Sadakane K, Lam T-W. MEGAHIT: an ultra-fast single-node solution for large and complex metagenomics assembly via succinct de Bruijn graph. Bioinformatics. 2015;31:1674–6.

99. Bushnell B, Rood J, Singer E. BBMerge – Accurate paired shotgun read merging via overlap. PLOS ONE. 2017. p. e0185056. Available from: http://dx.doi.org/10.1371/journal.pone.0185056

100. Peng Y, Leung HCM, Yiu SM, Chin FYL. IDBA-UD: a de novo assembler for single-cell and metagenomic sequencing data with highly uneven depth. Bioinformatics. 2012;28:1420–8.

101. Huang Y, Niu B, Gao Y, Fu L, Li W. CD-HIT Suite: a web server for clustering and comparing biological sequences. Bioinformatics. 2010;26:680–2.

102. BamM - Working with the BAM. Available from: http://ecogenomics.github.io/BamM/

103. Quinlan AR. BEDTools: the Swiss-army tool for genome feature analysis. Curr Protoc Bioinformatics. Wiley Online Library; 2014;47:11–2.

104. Paez-Espino D, Chen I-MA, Palaniappan K, Ratner A, Chu K, Szeto E, et al. IMG/VR: a database of cultured and uncultured DNA Viruses and retroviruses. Nucleic Acids Res. 2017;45:D457–65.

105. Roux S, Trubl G, Goudeau D, Nath N, Couradeau E, Ahlgren NA, et al. Optimizing de novo genome assembly from PCR-amplified metagenomes. PeerJ. 2019;7:e6902.

106. Ren J, Ahlgren NA, Lu YY, Fuhrman JA, Sun F. VirFinder: a novel k-mer based tool for identifying viral sequences from assembled metagenomic data. Microbiome. 2017;5:69.

107. Bushnell B. BBMap: a fast, accurate, splice-aware aligner. Lawrence Berkeley National Lab.(LBNL), Berkeley, CA (United States); 2014. Available from: https://www.osti.gov/biblio/1241166

108. Kang DD, Froula J, Egan R, Wang Z. MetaBAT, an efficient tool for accurately reconstructing single genomes from complex microbial communities. PeerJ. 2015. p. e1165. Available from: http://dx.doi.org/10.7717/peerj.1165

109. Parks DH, Imelfort M, Skennerton CT, Hugenholtz P, Tyson GW. CheckM: assessing the quality of microbial genomes recovered from isolates, single cells, and metagenomes. Genome Res. 2015;25:1043–55.

110. Sorensen JW, Dunivin TK, Tobin TC, Shade A. Ecological selection for small microbial genomes along a temperate-to-thermal soil gradient. Nat Microbiol. 2019;4:55–61.

111. Johnston ER, Hatt JK, He Z, Wu L, Guo X, Luo Y, et al. Responses of tundra soil microbial communities to half a decade of experimental warming at two critical depths. Proc Natl Acad Sci U S A. 2019;116:15096–105.

112. Wu Y-W, Simmons BA, Singer SW. MaxBin 2.0: an automated binning algorithm to recover genomes from multiple metagenomic datasets. Bioinformatics. 2016;32:605–7.

113. Langmead B, Salzberg SL. Fast gapped-read alignment with Bowtie 2. Nat Methods. 2012;9:357–9.

114. Li H, Handsaker B, Wysoker A, Fennell T, Ruan J, Homer N, et al. The Sequence Alignment/Map format and SAMtools. Bioinformatics. 2009;25:2078–9.

115. Parks DH, Rinke C, Chuvochina M, Chaumeil P-A, Woodcroft BJ, Evans PN, et al. Recovery of nearly 8,000 metagenome-assembled genomes substantially expands the tree of life. Nat Microbiol. 2017;2:1533–42.

116. Olm MR, Brown CT, Brooks B, Banfield JF. dRep: a tool for fast and accurate genomic comparisons that enables improved genome recovery from metagenomes through de-replication. ISME J. 2017;11:2864–8.

117. Chaumeil P-A, Mussig AJ, Hugenholtz P, Parks DH. GTDB-Tk: a toolkit to classify genomes with the Genome Taxonomy Database. Bioinformatics. 2019; Available from: http://dx.doi.org/10.1093/bioinformatics/btz848

118. Parks DH, Chuvochina M, Waite DW, Rinke C, Skarshewski A, Chaumeil P-A, et al. A standardized bacterial taxonomy based on genome phylogeny substantially revises the tree of life. Nat Biotechnol. 2018;36:996–1004.

119. Hyatt D, Chen G-L, Locascio PF, Land ML, Larimer FW, Hauser LJ. Prodigal: prokaryotic gene recognition and translation initiation site identification. BMC Bioinformatics. 2010;11:119.

120. Eddy SR. Accelerated Profile HMM Searches. PLoS Comput Biol. 2011;7:e1002195.

121. Zinke LA, Evans PN, Schroeder A, Parks DH. Evidence for non-methanogenic metabolisms in globally distributed archaeal clades basal to the Methanomassiliicoccales. bioRxiv. biorxiv.org; 2020; Available from: https://www.biorxiv.org/content/10.1101/2020.03.09.984617v1.abstract

122. Burstein D, Harrington LB, Strutt SC, Probst AJ, Anantharaman K, Thomas BC, et al. New CRISPR-Cas systems from uncultivated microbes. Nature. 2017;542:237–41.

123. Kumar S, Stecher G, Li M, Knyaz C, Tamura K. MEGA X: Molecular Evolutionary Genetics Analysis across Computing Platforms. Mol Biol Evol. 2018;35:1547–9.

124. Hug LA, Baker BJ, Anantharaman K, Brown CT, Probst AJ, Castelle CJ, et al. A new view of the tree of life. Nat Microbiol. 2016;1:16048.

125. Oksanen J, Blanchet FG, Friendly M, Kindt R, Legendre P, McGlinn D, et al. vegan: Community Ecology Package. R package version 2.4-3. Vienna: R Foundation for Statistical Computing. 2016;

126. Love MI, Huber W, Anders S. Moderated estimation of fold change and dispersion for RNA-seq data with DESeq2. Genome Biol. 2014;15:550.

