## supplemental materals, methods, discussion and figures for "Minnesota peat viromes reveal terrestrial and aquatic niche partitioning for local and global viral populations"

### Supplemental discussion

#### *Auxiliary metabolic gene analysis*

In order to identify putative viral auxiliary metabolic genes (AMGs) that may have a role in host metabolism during the infection cycle, and so that we could compare the output of two recently developed software tools for AMG prediction, we used VIBRANT [1] and DRAM-v [2] to recover putative AMGs from the SPRUCE vOTUs. We ran VIBRANT [1] on the 4,326 SPRUCE vOTUs that were predicted by either VirSorter [3] or DeepVirFinder [4], resulting in 2,802 vOTUs used for the VIBRANT analysis. Because DRAM-v requires VirSorter output, we re-ran the 4,326 SPRUCE vOTUs through VirSorter, resulting in 3,870 vOTUs, of which we only retained 2,645 for DRAM-v that were also recovered by VIBRANT. For DRAM-v output, only putative AMGs with an auxiliary score < 4 were retained, and no viral flag (F), transposon flag (T), viral-like peptidase (P) or attachment flag (A) could be present, and putative AMGs that did not have a gene ID or a gene description were also discarded.

We manually scanned the annotation output from both programs to give an indication as to whether the predicted AMGs were likely to be in *bona fide* viral regions of contigs. In particular, we were looking for: 1) the presence of viral genes upstream and downstream of the putative AMG, 2) putative AMG not in a genomic region with other host-like genes that could represent an operon (potentially indicative of miscalled ends of proviral sequences in the original vOTU sequence), and 3) an overall contig annotation consistent with a viral origin, such as the presence of viral hallmark genes, as a small number of false-positive vOTUs are known to be called by both VirSorter and DeepVirFinder [1,3,4]. Although DRAM-v performs an automated version of some of these manual investigation steps, we found that it was still useful to manually scan the DRAM-v output. It was not practical to manually curate the entire dataset, so we performed annotation spot-checks at random to get an overall sense of the quality of the output, which we describe below.

In total, 837 putative AMGs were predicted by VIBRANT (Table S6, Table S10) and 456 by DRAM-v (Table S11) prior to manual investigation. The VIBRANT output file with the predicted metabolic pathway(s) for each putative AMG revealed the presence of some genes in more than one pathway, resulting in duplicate entries of some genes as AMGs. Of these, we manually investigated putative AMGs that were involved in highly specific metabolic pathways that would not be expected in viruses, such as xenobiotic biodegradation and metabolism and biosynthesis of secondary metabolites like antibiotics, to see whether additional pathway results might be more intuitive than these putative assignments. In most of these cases, an alternative assigned pathway and/or gene description had been seen in viruses before (*e.g.*, nucleotide metabolism, consistent with viral replication needs, or carbon metabolism, consistent with known AMGs [5–7]), so the more specific pathways were removed from further consideration, and the more virus-consistent pathway calls were retained. VIBRANT and DRAM-v also recovered 50 and 48 putative AMGs predicted to be involved in nucleotide metabolism, respectively, which we removed from downstream consideration. We also discarded putative AMGs predicted to be glycosyl transferases, because some viruses can encode their own glycosyl transferases [8–10]. After these curation steps, 787 putative AMGs were retained from VIBRANT and 255 from DRAM-v, or 67% and 56% of the originally identified putative AMGs, respectively.

VIBRANT placed most of the remaining putative AMGs in the metabolism of cofactors and vitamins category (n=217). Upon closer examination, we found that some of these genes were glycosyl transferases (n=37), adenylyltransferases (n=26), or methyltransferases (n=10), which have been described previously in viral genomes [11,12, 13]. In fact, the general category of “methyltransferase” was the single most common annotation in hypersaline lake viromes [11]. DRAM-v placed most of its remaining putative AMGs in the ‘Organic Nitrogen’ category (n=89), but upon closer inspection, many of these putative AMGs were either peptidases (consistent with a viral origin [12]) or involved in amino acid metabolism. VIBRANT similarly classified 151 putative AMGs in the ‘Amino acid metabolism’ class, where the most common enzyme was methyltransferase (n=93), which, again, is a common annotation for viruses [11,13,14]. Thus, we would conservatively exclude these putative AMGs from consideration, barring further manual investigation of their true predicted functions and genomic contexts.

The second-largest putative AMG metabolic class for both VIBRANT and DRAM-v was carbon metabolism. VIBRANT classified a total of 212 putative AMGs in this category, but also 75 putative AMGs were assigned to the glycan metabolism category, which is a more specific kind of carbon metabolism, for a total of 287 putative carbon-cycling AMGs. DRAM-v identified 88 putative AMGs in the carbon metabolism category. These types of AMGs have been relatively well studied in marine systems [6,15–17] and have also been found in soil viruses [5,18], consistent with previously proposed direct viral roles in soil carbon cycling. Other putative AMGs that VIBRANT found were placed in pathways for the metabolism of terpenoids and polyketide sugars (which could also be classified as carbon-cycling, since these are large carbon molecules), energy metabolism, lipid metabolism, sulfur metabolism, and biosynthesis of other secondary metabolites. Other putative AMGs identified by DRAM-v were mostly transport proteins and ribosomal proteins. Transport proteins have been recovered as putative AMGs before [16,17], but ribosomal proteins have been shown to be present in some viral genomes for interacting with the host cell’s translation machinery [19], so, conservatively, we do not consider them to be likely AMGs.

Based on our experience with both programs, we recommend manual curation of the output to improve confidence in AMG identification. Given that AMGs were not the primary focus of this work, a comprehensive, labor-intensive manual curation effort was beyond the scope of this study, but we do recommend manual curation for future studies that rely more heavily on high-quality AMG predictions, and/or for specific AMGs of interest in a given dataset. Our results are consistent with the logical assumption that these tools perform best on well-annotated, *bona fide* vOTUs and perhaps less well on poorly annotated and/or misidentified viral genomic regions in the input data. For example, manual scanning of the output from both programs revealed some putative host operons identified as AMGs (*i.e.*, multiple cellular metabolism-like genes in a row, which would be less consistent with a true viral origin and perhaps more likely to be an error in the initial virus prediction, *e.g.*, a miscalled end of a proviral contig that read into the host genome). Thus, interestingly, these tools might also be useful for assisting with manual curation of viral datasets by allowing researchers to hone in on "putative AMGs" that might actually be non-viral genomic regions that passed through viral detection software. In summary, although these tools are certainly helpful for mining AMGs from metagenomic data, manual curation of AMG predictions is particularly recommended for total metagenomic datasets, which may include large numbers of novel vOTUs that could contain contaminating genomic regions derived from cellular genomes.

### Supplemental methods

#### vOTUs from the Alaskan peat dataset

##### *Sample collection*

Three peat soil cores were collected in June 2013 from the active layer of a *Sphagnum*-dominated thermokarst bog in a zone of discontinuous permafrost as part of the Alaska Peatland

Experiment (APEX) program, which is part of the Bonanza Creek LTER southwest of Fairbanks, AK (64.70 °N, -148.3 °W). Moss and peat were removed until the top of the water table was reached, and the top 40 cm below the water table were collected using a 6.3 cm diameter barrel corer. Mason jars were filled with porewater from the core hole, and the peat cores were stored in these mason jars.

*Incubations*

Jars were sealed and shipped on ice to the U.S. Geological Survey (Menlo Park, CA, USA). Samples were homogenized in an anaerobic glovebox at 4 °C. From each core, a total of four homogenized samples (2 g of soil wet weight) were collected in triplicate, creating 12 incubations. Samples were pressed to remove porewater and put into Wheaton serum vials that were sealed with blue butyl rubber stoppers. $O_2$ was removed from the glovebox atmosphere by vacuuming and filling with $N_2$ (10 in Hg/5 psi) 10 times in a cold room (4 °C) to purge the headspaces of the vials. Soil porewater was freeze-dried, and remaining particulates were rehydrated with either heavy water (97 atom% $H_2{}^{18}O$, Cambridge Isotope Laboratories) or natural abundance water (control) and then sparged with $N_2$ to remove $O_2$, in order to create synthetic porewater. The samples were put in a glycerol bath at 4 °C, and the temperature was reduced slowly and steadily to -1.5 °C over 48 h. Samples were maintained at -1.5 °C. Per treatment, half of the samples were destructively sampled (in triplicate) at 184 d and 370 d. The samples were snap-frozen in liquid $N_2$ and stored at -80 °C until DNA extraction.

*DNA extraction, DNA isotopic fractionation, library construction, and sequencing*

DNA was extracted from 0.5 g wet soil per sample. Soil was added to lysing matrix E tubes (MP Biomedicals, Burlingame, CA), and 100 mL 1x TE (pH 7.5), 150 mL $PO_4^{3-}$ buffer (0.2 M in 1 M NaCl), and 100 µl 10% SDS were added. Tubes were vortexed for 30 s and briefly centrifuged at low speed to consolidate the sample in the bottom of the tubes. 600 µl of phenol:chloroform:isoamyl alcohol (PCI, 25:24:1) was added, and then tubes were vortexed and incubated at 65°C for 10 min. Tubes were then centrifuged for 5 min at 10,000 x g, and the supernatant was put into a new tube. Re-extraction was done on the lysing matrix E tubes using 220 mL 1x TE and 80 µl $PO_4^{3-}$ buffer. The supernatants of these extractions were combined in a new 2 ml tube. 550 µl of PCI was added, homogenized, and centrifuged (10,000 x g, 5 min). The aqueous top layer was transferred to a new 2 ml tube, and 900 µl of chloroform:isoamyl alcohol (CI, 24:1) was added, mixed, and centrifuged at 10,000 x g for 5 min. Supernatant was put into a new 2 mL tube, and 850 µl CI was added, mixed, and centrifuged at 10,000 x g for 5 min, and then the supernatant was added to a new 1.7 ml tube. RNase was added (6.44 µl, 10 mg/ml), mixed, and incubated at 50 °C for 10 min. Then 244 µl 10 M ammonium acetate was added, mixed, and incubated (4 °C, 2 h). Tubes were then centrifuged at 16,100 x g for 15 min. Supernatant was put into a new tube, and isopropanol (670 µl) was added, mixed, and centrifuged (16,100 x g, 20 min). The supernatant was then discarded, and the DNA pellet was dried in a PCR hood for 15 min. The DNA pellet was resuspended in 30 µl 1x TE, and DNA was stored at -80 °C.

CsCl density gradient ultracentrifugation was done on the extracted DNA in order to separate DNA and $^{18}O$-enriched DNA. DNA from five density fractions was collected, and DNA from the two heaviest fractions (medium-heavy [MH; 1.717–1.725 g/ml] and heavy [H; 1.725–1.750 g/ml]) was collected and sheared to 500 bp using the Covaris E210, then a SPRI bead cleanup was done using Ampure XP SPRI and quality-controlled using a Bioanalyzer chip. Sheared DNA was then end-repaired, A-tailed, and adapter-ligated. A total of 24 metagenomic libraries were constructed and sequenced. Sequencing of these libraries was done on an Illumina HiSeq 2500 and was successful for 23 samples, yielding a total of 302 Gbp of sequencing data.

*Bioinformatics processing and vOTU recovery*

Bbduk (v38.56) was used for trimming raw reads (151 nt, PE) (ftl=10 ktrim=r k=23 mink=11 hdist=1 tpe tbo minlen=50), PhiX and Illumina adapter/barcode removal (k=31 hdist=1 minlen=50), and quality trimming (qtrim=r trimq=10 minlen=50). Bioinformatics specific for viral recovery were used to increase the number of viral contigs detected in these datasets [5,20]. Reads were assembled into contigs using SPAdes v3.11.1 (--only-assembler --phred-offset 33 --meta -k 25,55,95 --12). VirSorter (virome decontamination mode; [3]) and DeepVirFinder (DVF; [4]) were used to detect viral contigs. Only contigs ≥ 10 kbp from VirSorter categories 1 and 2 or with DeepVirFinder scores ≥ 0.9 and $p < 0.05$ were retained. All resulting viral contigs were clustered into vOTUs using nucmer [22] at 95% average nucleotide identify (ANI) across 85% of the shorter contig [21].

*Availability of Alaskan peat metagenomic data*

The 23 Arctic metagenomes were deposited to NCBI under BioProject identifier PRJNA634918.

### vOTUs from the Puerto Rican forest soil dataset

*Sample collection and metagenomic data generation*

Soils collected from the Luquillo Experimental Forest (LEF), El Verde Field Station, Puerto Rico, USA (18°18′ N; 65°50 ′W) [23] are clay and Fe oxide-rich volcanoclastic oxisols with 5.8% carbon and pH of ~5. Homogenized soil was amended with $^{13}C$ ryegrass (99 atm %, IsoLife) and incubated under four redox treatments-- static oxic, low frequency redox fluctuation (8-days oxic, 4-days anoxic), high frequency redox fluctuation (4-days oxic, 4-days anoxic), and static anoxic conditions, to assess effects of redox periodicity on the soil microbial community and C transformations [23–25]. From soils harvested at 0 and 44 weeks, DNA was extracted (using a modified Griffith's protocol using 0.25 g/ microcosm, extracted in triplicate and then pooled for downstream sequencing) [26] and fractionated as described above for the Alaskan peat samples. In total, 95 metagenomes (85 stable isotope probing (SIP) fractions from the 'heavy' density fractions (> 1.73 g/ml) and 10 non-labeled bulk soils) were sequenced at the Joint Genome Institute, with the Illumina NovaSeq Low Input Protocol (DNA) as 270 bp fragments.

*Bioinformatics*

Reads were processed using the JGI QC pipeline (https://jgi.doe.gov/data-and-tools/bbtools/bb-tools-user-guide/data-preprocessing/ , accessed January 6, 2019) and were assembled into contigs using SPAdes v3.11.1 (--only-assembler --phred-offset 33 --meta -k 25,55,95 --12)[27]. Viral contigs were detected with VirSorter[3] from categories 1, 2, 4, and 5 and DeepVirFinder[4], score > 0.9 and p value < 0.05. Viral contigs were clustered into vOTUs based on 95% nucleotide similarity and 85% alignment fraction.

*Availability of metagenomic data*

NCBI accession numbers of the 95 metagenomes can be found in table S14.

1. Kieft K, Zhou Z, Anantharaman K. VIBRANT: automated recovery, annotation and curation of microbial viruses, and evaluation of viral community function from genomic sequences. Microbiome. 2020;8:90.

2. Shaffer M, Borton MA, McGivern BB, Zayed AA, La Rosa SL, Solden LM, et al. DRAM for distilling microbial metabolism to automate the curation of microbiome function. Nucleic Acids Res . 2020; Available from: http://dx.doi.org/10.1093/nar/gkaa621

3. Roux S, Enault F, Hurwitz BL, Sullivan MB. VirSorter: mining viral signal from microbial genomic data. PeerJ. 2015;3:e985.

4. Ren J, Song K, Deng C, Ahlgren NA, Fuhrman JA, Li Y, et al. Identifying viruses from metagenomic data using deep learning. Quantitative Biology. 2020;8:64–77.

5. Trubl G, Hyman P, Roux S, Abedon ST. Coming-of-Age Characterization of Soil Viruses: A User's Guide to Virus Isolation, Detection within Metagenomes, and Viromics . Soil Systems. 2020. p. 23. Available from: http://dx.doi.org/10.3390/soilsystems4020023

6. Hurwitz BL, U'Ren JM. Viral metabolic reprogramming in marine ecosystems. Curr Opin Microbiol. 2016;31:161–8.

7. Crummett LT, Puxty RJ, Weihe C, Marston MF, Martiny JBH. The genomic content and context of auxiliary metabolic genes in marine cyanomyoviruses. Virology. 2016;499:219–29.

8. Xiang Y, Baxa U, Zhang Y, Steven AC, Lewis GL, Van Etten JL, et al. Crystal structure of a virus-encoded putative glycosyltransferase. J Virol. 2010;84:12265–73.

9. Larson ET, Reiter D, Young M, Lawrence CM. Structure of A197 from Sulfolobus turreted icosahedral virus: a crenarchaeal viral glycosyltransferase exhibiting the GT-A fold. J Virol. 2006;80:7636–44.

10. Markine-Goriaynoff N, Gillet L, Van Etten JL, Korres H, Verma N, Vanderplasschen A. Glycosyltransferases encoded by viruses. J Gen Virol. 2004;85:2741–54.

11. Emerson JB, Thomas BC, Andrade K, Heidelberg KB, Banfield JF. New approaches indicate constant viral diversity despite shifts in assemblage structure in an Australian hypersaline lake. Appl Environ Microbiol. 2013;79:6755–64.

12. James MNG. The peptidases from fungi and viruses. Biol Chem. 2006;387:1023–9.

13. Jeudy S, Rigou S, Alempic J-M, Claverie J-M, Abergel C, Legendre M. The DNA methylation landscape of giant viruses. Nat Commun. 2020;11:2657.

14. Coutard B, Barral K, Lichière J, Selisko B, Martin B, Aouadi W, et al. Zika Virus Methyltransferase: Structure and Functions for Drug Design Perspectives. J Virol . 2017;91. Available from: http://dx.doi.org/10.1128/JVI.02202-16

15. Hurwitz BL, Hallam SJ, Sullivan MB. Metabolic reprogramming by viruses in the sunlit and dark ocean. Genome Biol. 2013;14:R123.

16. Warwick-Dugdale J, Buchholz HH, Allen MJ, Temperton B. Host-hijacking and planktonic piracy: how phages command the microbial high seas. Virol J. 2019;16:15.

17. Brum JR, Sullivan MB. Rising to the challenge: accelerated pace of discovery transforms marine virology. Nat Rev Microbiol. 2015;13:147–59.

18. Emerson JB, Roux S, Brum JR, Bolduc B, Woodcroft BJ, Jang HB, et al. Host-linked soil viral ecology along a permafrost thaw gradient. Nature Microbiology. Nature Publishing Group; 2018;3:870–80.

19. Mizuno CM, Guyomar C, Roux S, Lavigne R, Rodriguez-Valera F, Sullivan MB, et al. Numerous cultivated and uncultivated viruses encode ribosomal proteins. Nat Commun. 2019;10:752.

20. Roux S, Trubl G, Goudeau D, Nath N, Couradeau E, Ahlgren NA, et al. Optimizing de novo genome assembly from PCR-amplified metagenomes. PeerJ. 2019;7:e6902.

21. Roux S, Adriaenssens EM, Dutilh BE, Koonin EV, Kropinski AM, Krupovic M, et al. Minimum Information about an Uncultivated Virus Genome (MIUViG). Nat Biotechnol. 2019;37:29–37.

22. Delcher AL, Salzberg SL, Phillippy AM. Using MUMmer to identify similar regions in large sequence sets. Curr Protoc Bioinformatics. 2003;Chapter 10:Unit 10.3.

23. Bhattacharyya A, Campbell AN, Tfaily MM, Lin Y, Kukkadapu RK, Silver WL, et al. Redox Fluctuations Control the Coupled Cycling of Iron and Carbon in Tropical Forest Soils. Environ Sci Technol. 2018;52:14129–39.

24. Lin Y, Bhattacharyya A, Campbell AN, Nico PS, Pett-Ridge J, Silver WL. Phosphorus Fractionation Responds to Dynamic Redox Conditions in a Humid Tropical Forest Soil. Journal of Geophysical Research: Biogeosciences. John Wiley & Sons, Ltd; 2018;123:3016–27.

25. Lin Y, Campbell AN, Bhattacharyya A, DiDonato N, Thompson AM, Tfaily MM, et al. Differential effects of redox conditions on the decomposition of litter and soil organic matter. Biogeosci Discuss. Copernicus GmbH; 2020;1–25.

26. Griffiths RI, Whiteley AS, O’Donnell AG, Bailey MJ. Rapid method for coextraction of DNA and RNA from natural environments for analysis of ribosomal DNA- and rRNA-based microbial community composition. Appl Environ Microbiol. 2000;66:5488–91.

27. Bankevich A, Nurk S, Antipov D, Gurevich AA, Dvorkin M, Kulikov AS, et al. SPAdes: a new genome assembly algorithm and its applications to single-cell sequencing. J Comput Biol. 2012;19:455–77.

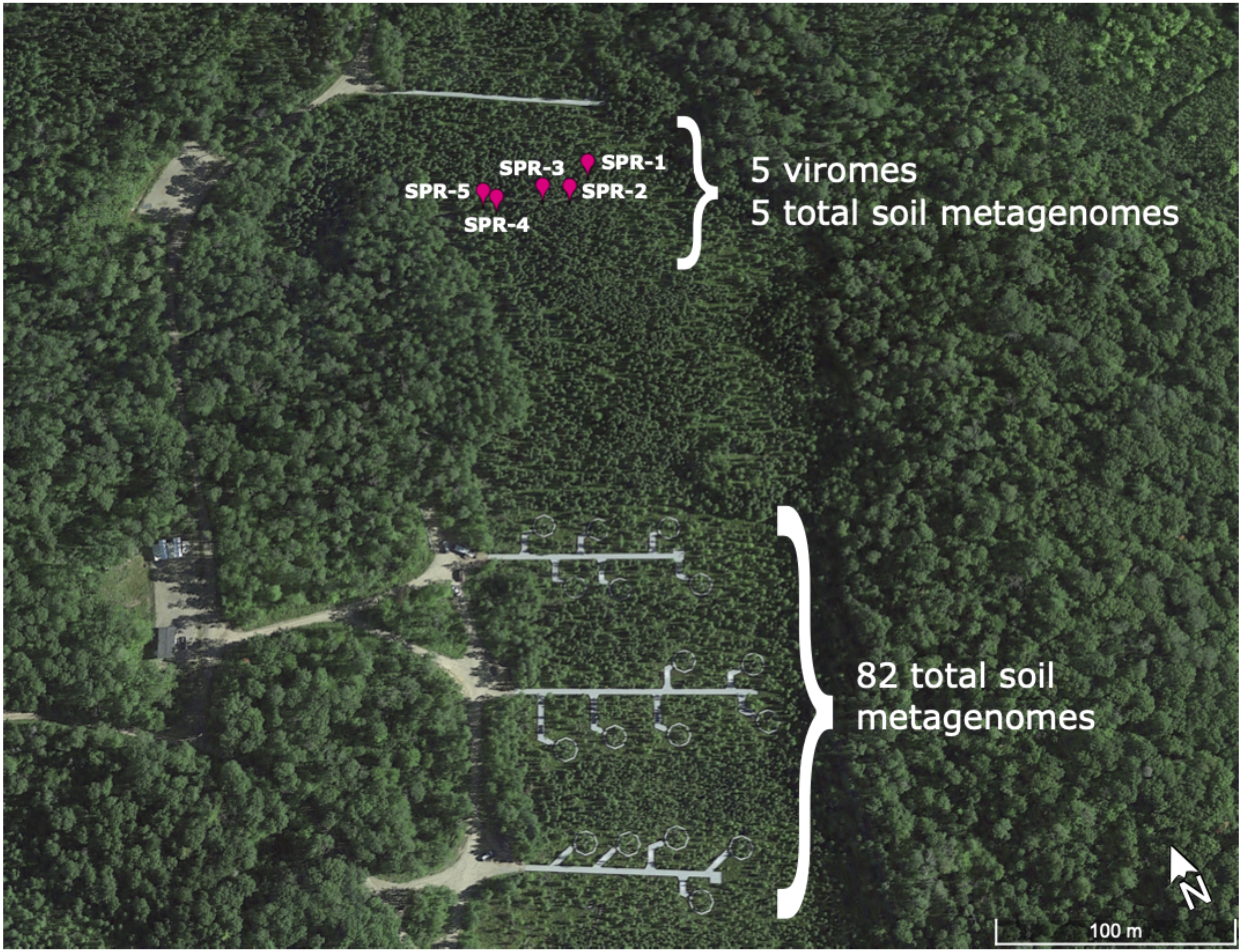


**Supplementary figure 1**: **Sampling locations for all SPRUCE samples**. Sampling locations within the S1 Bog at the Marcell Experimental Forest in Northern Minnesota, USA, including the five transect samples and the samples from the SPRUCE experimental chambers. Numbers next to the brackets show how many and what kinds of metagenomes were derived from each part of the bog.

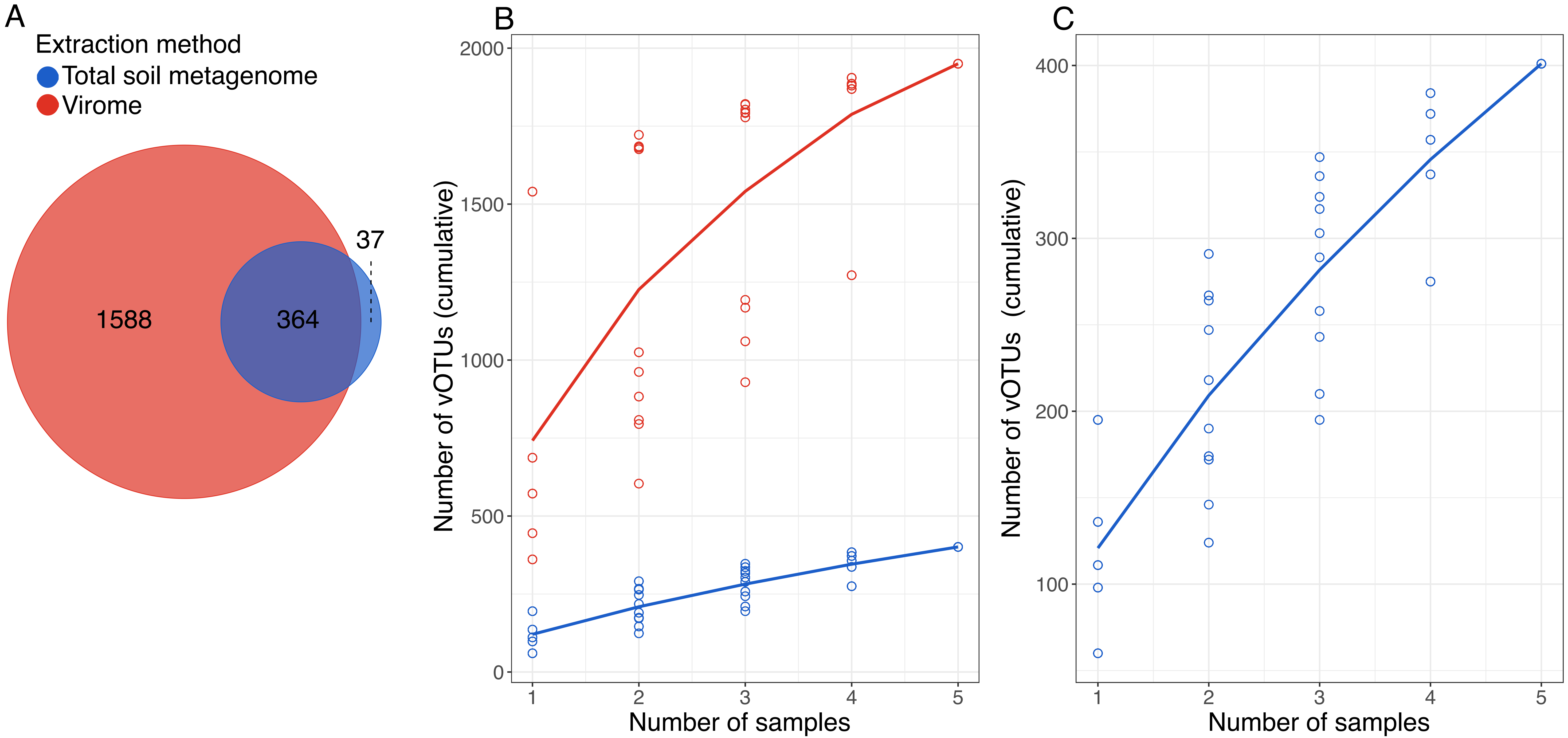


**Supplementary figure 2**: **Comparison of vOTU recovery from five paired viromes and total soil metagenomes from the SPRUCE transect. A**: Distribution of vOTUs recovered by each of the two extraction methods, based on read mapping to the PIGEON database, including all vOTUs recovered from SPRUCE. **B**: Accumulation curves of distinct vOTUs recovered as sampling increases for each extraction method; 100 permutations of sample order are depicted as open circles, and averages are shown as a line. **C**: Similar to panel B, but only the accumulation curve of distinct vOTUs recovered from total soil metagenomes is shown, with a smaller y-axis maximum to better show the trend.

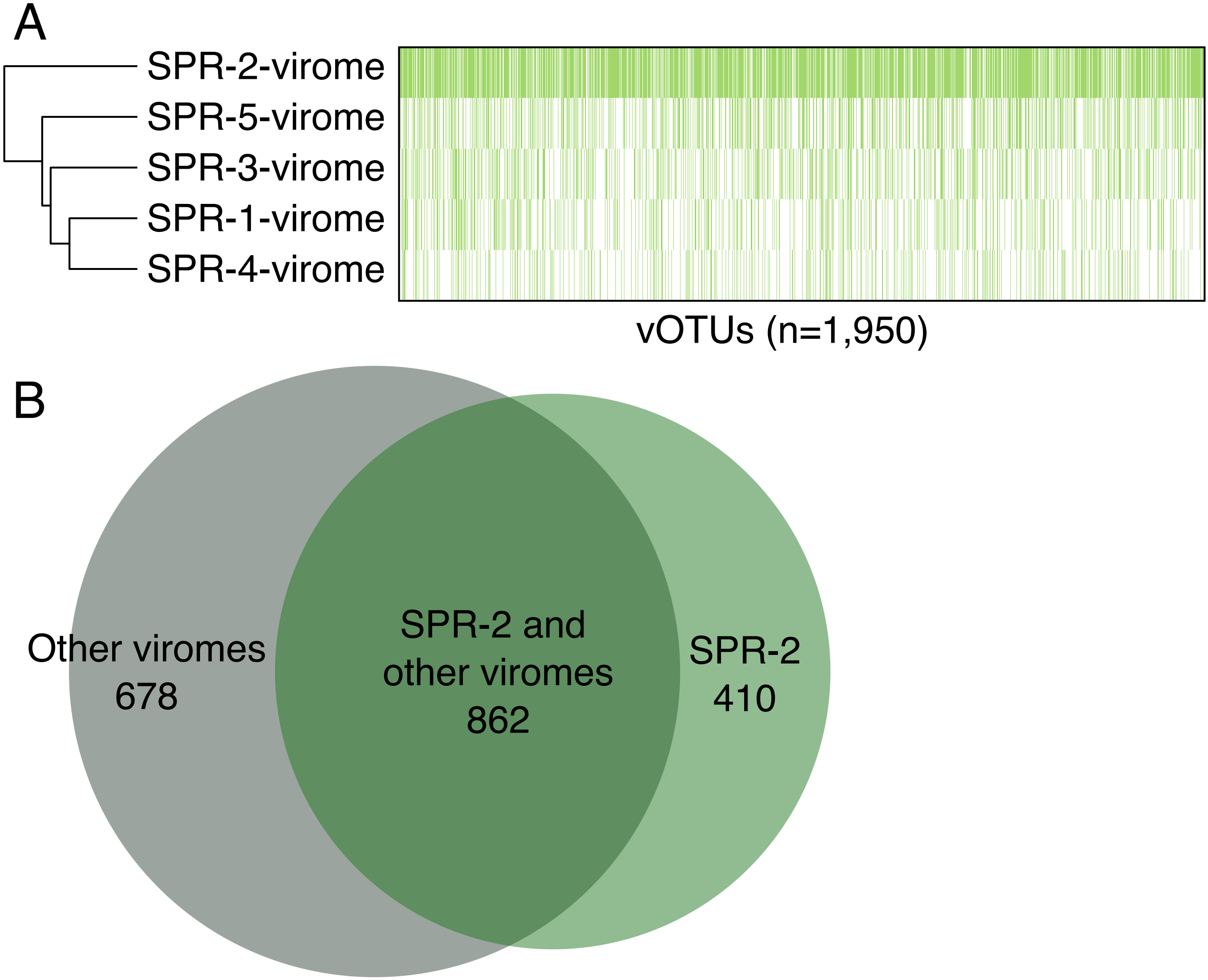


**Supplementary figure 3: Comparison of the five viromes from the transect. A:** Dendrogram depicting sample similarity according to viral community composition (left) and heatmap (right) of vOTUs detected (green = detected, white = not detected) in the five SPRUCE transect viromes.
**B:** Comparison of vOTU recovery from the SPRUCE-2 sample compared to the four other virome samples.
